# Grey matter embedding of the cholinergic system

**DOI:** 10.64898/2025.12.02.690396

**Authors:** Alexander Weuthen, Sidhant Chopra, Dario Lucas Bergmann, Meng Li, Bianca Besteher, Martin Walter

## Abstract

Integrity of acetylcholine-producing basal forebrain nuclei Ch1-4 is crucial for neurocognitive functioning and grey matter decline represents a hallmark of neuropsychiatric disorders, including Alzheimer’s disease, Parkinson’s disease and schizophrenia. Approaches that link brain-wide grey matter atrophy with diminished cholinergic innervation from Ch1-4 projections are required for neuroimaging of disease stages and pharmacological decisions addressing cholinergic deficits. The current study assessed human magnetic resonance imaging-based markers for cholinergic nucleus subregions and synaptic targets in healthy exploration (HCP, *n* = 1113) and replication (IXI, *n* = 587) cohorts. Ch1-3 and Ch4 were delineated at 50% cytoarchitectonic probability to determine their grey matter coupling patterns using structural covariance analyses. Spatial colocalization with vesicular acetylcholine transporter, M1-muscarinic and α4β2-nicotinic receptor density quantified the correspondence with synaptic cholinergic markers. All individual-level cholinergic nucleus and target region indices showed good test-retest reliability (*R* ≥ 0.80) when restricted to voxels with at least 10% grey matter probability. Spatial colocalization with presynaptic acetylcholine transporters emphasized a cholinergic nature of Ch1-4 grey matter and its coupling with corticopetal projections such as amygdala, insula and cingulate cortices. Subcortical gradients of M1-muscarinic receptor density and the Ch1-3 structural covariance map, however, dominated in locations of striatal cholinergic interneurons, such as nucleus accumbens, caudate nucleus and putamen. Ch4 covaried with medial thalamus and hippocampus, while both subregions’ structural covariance networks converged in a posterolateral Ch4 extensions, potentially resembling subputaminal nucleus of Ayala. The results emphasize Ch1-4 morphometry as marker for acetylcholine-driven brain regions and describe neuroimaging strategies to indicate synaptic cholinergic involvement.

**Significance Statement:** Previous studies have suggested that grey matter alterations in basal forebrain Ch1-4 nuclei indicate cholinergic deficits in neurodegenerative and psychiatric pathologies. However, the use of Ch1-4 volume as a marker of cholinergic system integrity has yet to be validated. Additionally, there is a lack of non-invasive procedures to quantify integrity of regions innervated by its cholinergic projections. Our results suggest that grey matter coupling with Ch1-4 is a sensitive cholinergic system marker, mimicking the density distribution of synaptic cholinergic innervation. Furthermore, we introduce grey matter-based target region indices, evaluate their robustness in individual participants and describe regional contributions to M1-muscarinic, α4β2-nicotinic and vesicular acetylcholine transporter gradients. The study advances our understanding of cholinergic circuits *in vivo* using magnetic resonance imaging.

## Introduction

The neurotransmitter acetylcholine modulates central and peripheral nervous system functioning via nicotinic (nAChR) and muscarinic (mAChR) receptors (Dale, 1914; Loewi, 1922). Cholinergic actions on nAChR and mAChR affect brain development through neurite extension and synaptogenesis (Bruel-Jungerman et al., 2011; Roqué et al., 2023), while axonally back-propagating nerve growth factor regulates the functional maintenance and survival of cholinergic neurons (see Cuello and Do Carmo, 2025 for a review). As such, cholinergic nuclei and their synaptic target regions are intrinsically linked throughout development and into adulthood. The basal forebrain contains magnocellular acetylcholine-projecting neuronal cell populations (Ch1-4) which predominantly target hippocampus from Ch1 in medial septum and Ch2 in vertical limb of diagonal band of Broca, olfactory bulbus from Ch3 in horizontal diagonal band, as well as amygdala and cerebral cortex from Ch4 in basal nucleus of Meynert subregions (Mesulam et al., 1983a). Experimental lesioning of rodent Ch1-4 neurons has been shown to reduce the dendritic arborization of layer V pyramidal cells and cortical thickness (Robertson et al., 1998; Hohmann, 2003). This suggests that synaptic cholinergic signalling influences grey matter morphology in its projection targets.

In humans, cytoarchitectonic studies found atrophy and reduced numbers of cholinergic neurons in Ch1-4 and indicated cholinergic deficits in neurodegenerative pathologies such as Alzheimer’s disease, Parkinson’s disease and dementia with Lewy bodies (Whitehouse et al., 1982; Mesulam and Geula, 1988; Mufson et al., 2003). With the availability of post-mortem cytoarchitectonic probability maps (Zaborszky et al., 2008), structural magnetic resonance imaging (MRI) studies have been able to non-invasively delineate the location of Ch1-4 and studies provided evidence for grey matter alterations in neurodegenerative pathologies (Kilimann et al., 2014; Schmitz and Spreng, 2016; Teipel et al., 2018; Schumacher et al., 2021; Crowley et al., 2024) and psychotic disorders (Avram et al., 2021; Schulz et al., 2023). Since cholinergic synapses are involved in grey matter maintenance (Robertson et al., 1998; Hohmann, 2003), a decline in Ch1-4 neurons likely accompanies grey matter atrophy in its neocortical, amygdalar and hippocampal projections. Therefore, the spatial distribution of widespread atrophy patterns in neurodegenerative conditions may at least partially be explained by and covary with Ch1-4 atrophy. More generally, structural covariance analyses seeded in Ch1-4 could capture spatial effects driven by cholinergic synapses in health and disease, considering structural maintenance and regional vulnerability in its cholinergic projection targets. Molecular validation could benefit from the successful development of positron emission tomography (PET) radioligands that selectively bind to acetylcholine’s synaptic targets. Previous PET studies have validated radioligand binding for the vesicular acetylcholine transporter (VAChT) (Petrou et al., 2014; Aghourian et al., 2017), M1-mAChR (Mogg et al., 2018; Naganawa et al., 2021) and α4β2-nAChR (Sabri et al., 2015; Hillmer et al., 2016), which enables spatial colocalization analyses with respective normative density topographies. Initial evidence revealed grey matter coupling between cholinergic nucleus and target regions in older adults with mild cognitive impairment (Cantero et al., 2016) and in patients with Alzheimer’s disease, where grey matter decline coupled to Ch1-4 spatially colocalized with VAChT density (Schmitz et al., 2018). VAChT density best resembles overall cholinergic innervation. Analyses of cholinergic receptor subtypes could delineate different cholinergic projection circuits and provide insight into the relative contributions of faster α4β2-nAChR ionotropic versus slower M1-mAChR metabotropic effects. Indexing explainable contributions of these receptors at the individual-participant level could provide neurobiology-informed quantification of cholinergic system status and potentially inform clinical indication of pharmacological interventions that selectively target these receptor subtypes.

The current study had two aims and analytical procedures (**Fig. 1**). First, we aimed to evaluate the external validity of grey matter volume of Ch1-4 and its subregions for synaptic cholinergic markers (**Fig. 1A**), which would highlight its potential as a marker of cholinergic system integrity. If Ch1-4 grey matter volume has a cholinergic basis, then grey matter throughout the brain should be coupled in regions with high cholinergic innervation, as inferred from colocalization with VAChT density. Based on post-mortem studies (Mesulam et al., 1983a), we expected structural covariance of Ch1-3 with hippocampus and of Ch4 with amygdala and neocortical regions. Second, we aimed to develop single-participant VAChT, M1-mAChR and α4β2-nAChR indices to complement the MRI-based evaluation of synaptic cholinergic target regions in addition to the assessment of cholinergic nucleus regions (**Fig. 1B**). Therefore, we quantified retest-reliability, overlaps and divergences of VAChT, M1-mAChR and α4β2-nAChR index measures embedded in the brain’s grey matter density distribution. We expected M1-mAChR and α4β2-nAChR indices to provide complementary insight into cholinergic system components, while being principally correlated with VAChT and Ch1-4 as more general cholinergic system markers.

**Figure 1.**
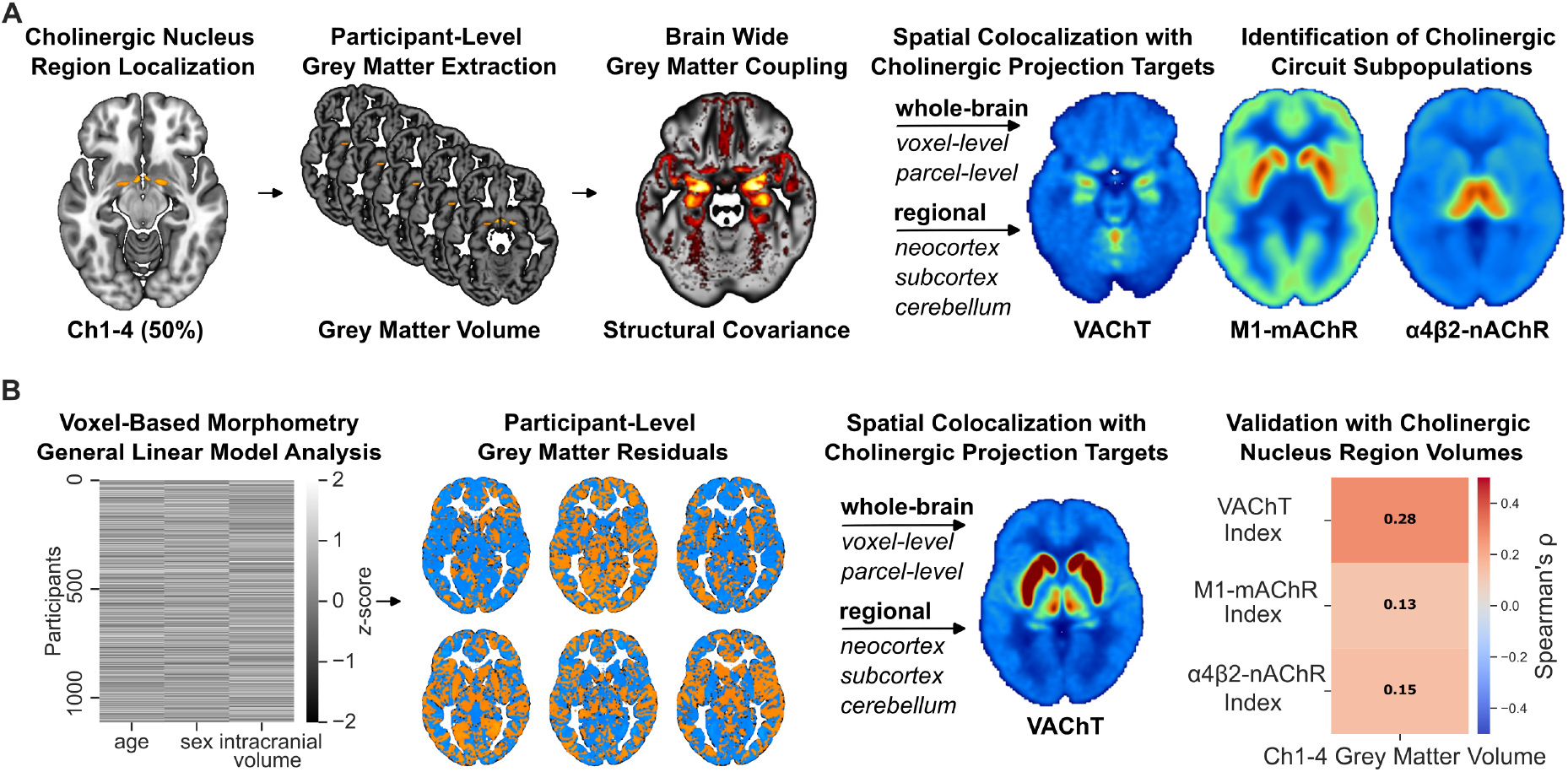
Methodological flowchart for reciprocal validation of cholinergic nucleus region and cholinergic projection target indices. **A**, Schematic representation of the structural covariance analyses of cholinergic nucleus region grey matter volume and assessment of molecular correspondence with cholinergic radioligand binding maps. **B**, Workflow for quantifying single-participant cholinergic target region indices based on participant’s brain-wide and regional grey matter expression.

## Materials and Methods

### Participants

The current study included participants from two study cohorts. The first cohort is based on the human connectome project (HCP) S1200 release (Van Essen et al., 2012) and contains 1113 participants (606 female, 507 male) with MRI scans within four age groups between 22-25, 26-30, 31-35 and 36+ years, as well as retest scans for 45 (31 female, 14 male) of the same participants. The second cohort is based on the Information eXtraction from Images (IXI, https://brain-development.org/ixi-dataset/) dataset and was used to test the overall replicability of the findings across samples. In the IXI dataset, MRI and relevant demographical data (age, sex) for 587 healthy participants (328 female, 259 male) within an age range between 18 and 86 years (*M* = 49.45, *SD* = 16.71) were available. Ethics approval and data sharing agreements for the studies protocols were obtained at respective study sites.

### MRI data acquisition

In the HCP dataset, MRI data were recorded with a customized Siemens 3T Skyra scanner at Washington University. T1-weighted MRI data were based on a magnetization prepared rapid gradient echo (MPRAGE) sequence (voxel size = 0.70 x 0.70 x 0.70 mm^3^, repetition time = 2.400 s, echo time = 2.14 ms, flip angle = 8°). The IXI dataset consisted of MRI images from different MRI scanning sites. T1-weighted structural MRI data were collected at three different hospitals using similar MPRAGE sequences (voxel size = 0.94 x 0.94 x 1.20 mm^3^, repetition time = 9.6/9.4/5.8 s, echo time = 4.6 ms, flip angle = 8°/8°/20°) in London, at 1) Hammersmith Hospital using a Philips 3T system, 2) Guy’s Hospital using a Philips 1.5T system and 3) Institute of Psychiatry using a GE 1.5T system.

### MRI data processing

MRI data from both datasets were available in a standardized neuroimaging format. To assess brain-wide voxel-based morphometry measures, the Computational Anatomy Toolbox (CAT12.9, Gaser et al., 2024) was used with MATLAB (version R2024a) to preprocess the structural MRI data applying standard options with shell script, and derived normalized grey matter probability maps for HCP with 0.7 x 0.7 x 0.7 mm^3^ voxel size and for IXI with optimized output resolution 1.5 x 1.5 x 1.5 mm^3^ to reduce resampling-induced inhomogeneities. Preprocessing involved tissue segmentation into grey matter, white matter and cerebrospinal fluid based on a unified segmentation approach and included registration to ICBM 2009c Nonlinear Asymmetric space (MNI152NLin2009cAsym) using a geodesic shooting algorithm. Quality control was based on the Image Quality Rating in the CAT12.9 metric (Gilmore et al., 2021), and showed that all images, both in HCP (*M* = 1.308, *SD* = 0.028, range 1.284 to 1.519) and IXI (*M* = 1.899, *SD* = 0.017, range 1.894 to 2.096) were rated as good or excellent.

### Cholinergic seed region definition

Delineation of Ch1-4 voxels was based on cytoarchitectonic probability maps (Zaborszky et al., 2008) released in the Julich-brain cytoarchitectonic atlas (version 3.1) (Amunts et al., 2020). Left and right regions were available for a combined Ch1-3 mask and for a non-overlapping Ch4 mask. Values in the mask corresponded to the proportion of individuals for which the voxel-based topography was cytoarchitectonically labelled as either Ch1-3 or Ch4 region. We chose a 50% probability threshold for Ch1-4 and its subregions probability maps for several reasons. First, we wanted to increase anatomical specificity for voxels containing cholinergic nuclei and to decrease overlap with adjacent structures such as cerebrospinal fluid, anterior commissure and amygdala. Additionally, a 50% probability threshold has been shown to correspond most closely with post-mortem histological volume for the Ch4 subregion (Wang et al., 2022). For the analysis on subregional effects, bilateral Ch1-3 and Ch4 masks were used respectively. For analyses on the whole Ch1-4, all subregional masks were combined.

### Seed region grey matter extraction

Cholinergic seed region masks were used to extract voxel-wise grey matter probability averages for Ch1-4_raw_, Ch1-3_raw_ and Ch4_raw_ for each participant. Subregional grey matter volumes were statistical orthogonalized, such that Ch1-3_ortho_ represented the residual grey matter volume after accounting for Ch4_raw_ and vice versa, as has been done in a previous study determining Ch1-3 and Ch4 functional connectivity patterns (Markello et al., 2018). Additional residuals independent from effects of age, sex and total intracranial volume were obtained from an ordinary least squares model, and then *z*-scored.

### Structural covariance analyses

Ch1-4_raw_, Ch1-3_ortho_ and Ch4_ortho_ structural covariance maps were assessed with separate univariate general linear model analyses. As described above, average grey matter probability of respective regions was extracted for each participant and defined as variable of interest. In the structural covariance analyses, we always included age, sex and total intracranial volume as additional nuisance regressors of no interest, as well as regressors for the three scanning sites of the IXI cohort. Analyses were restricted to voxels with at least 10% grey matter probability on each cohort’s average grey matter template and a voxel-wise false discovery rate threshold (*q* < .05) was used to account for multiple comparisons. Structural covariance analyses were first performed on the HCP cohort, and replicability of the results was later checked in the IXI cohort. While we focused on the analyses of the HCP cohort in the main results and figures, we show the structural covariance results of the IXI cohort at the same brain coordinates within in the Supplementary Material, and provide unthresholded maps on a public repository.

### Cholinergic target region definition

Density maps for cholinergic receptors (M1-mAChR, α4β2-nAChR) and transporters (VAChT) were based on publicly available group-average radioligand binding topographies of healthy control participants that were made public through the Neuromaps project (Markello et al., 2022). We downloaded slightly adapted maps, that included an MNI152NLin2009cAsym space transformation, brain extraction and scaling from 0.000001 to 1, available in the Nispace toolbox (Lotter and Dukart, 2024). The VAChT density map was based on binding of [^18^F]-FEOBV (Aghourian et al., 2017). This radioligand has been validated as a reliable marker of presynaptic VAChT density (Petrou et al., 2014). The M1-mAChR density map was based on binding of [^11^C]-LSN3172176 (Naganawa et al., 2021), which has been validated as a selective M1-mAChR receptor ligand (Mogg et al., 2018). The α4β2-nAChR density map was based on binding of [^18^F]-Flubatine (Hillmer et al., 2016), a validated and selective radioligand for α4β2-nAChR (Sabri et al., 2015). All of the above radiotracers have previously demonstrated high affinity, and there is limited evidence of off-target binding.

For the cerebral cortex, the Schaefer parcellation with 400 regions within volumetric MNI152NLin2009cAsym space was used (Schaefer et al., 2018), applying cytoarchitectonic annotations based on the Van-Economo-Koskinos atlas (Scholtens et al., 2018), as has been done in a study on brain stem and cortical associations (Hansen et al., 2024). For subcortical regions, the Tian parcellation version 2 with 16 left and 16 right regions was used to extract thalamus, amygdala, hippocampus, globus pallidus, caudate nucleus, putamen and nucleus accumbens subregions (Tian et al., 2020). For the cerebellum, the non-linear transformed parcellation with maximum probability maps of 50% in 1 mm MNI152NLin2009cAsym space was used (Diedrichsen et al., 2009), including 28 regions, while the vermis crus I region was not labelled in the cerebellar atlas and was therefore not part of the analyses. All maps were down-sampled to 3 x 3 x 3 mm^3^ resolution based on the MNI152NLin2009cAsym_res-3 template (Ciric et al., 2022) to account for the lower resolution of the PET data and to reduce excessive computation time in the correction for spatial autocorrelation. For each of the radioligand binding maps, 1000 surrogate null maps were computed based on the generative null model method available in the BrainSmash toolbox (Burt et al., 2020) using the implementation of the Neuromaps toolbox (Markello et al., 2022).

### Cholinergic indices extraction

For each participant a residualized grey matter map was derived from the group-level general linear model containing regressors for an intercept, age, sex and total intracranial volume effects. Extraction of regional grey matter probability and radioligand binding was restricted to voxels level having at least 10% grey matter probability in cohort’s grey matter average. Residual grey matter maps were assessed for their spatial colocalization with VAChT, M1-mAChR and α4β2-nAChR binding using Spearman correlation coefficients on a voxel-wise level within grey matter voxels. For the cerebral cortical parcellation, 400 regions were extracted for both the grey matter residuals and for the cholinergic target structure density and then correlated. This was repeated for the 32 subcortical and the 27 cerebellar regions, as well as for a whole brain parcellation including all 459 regions restricted to voxels in the 10% grey matter mask. In this way, each participant obtained five indices (whole brain grey matter voxel-wise, parcel-wise, cerebral cortex, subcortex, cerebellum) for each of the three cholinergic molecular target structures (VAChT, M1-mAChR, α4β2-nAChR), summing up to 15 cholinergic indices. For the assessment of intercorrelations between cholinergic indices and with regional grey matter measures, a Bonferroni-correction was applied to the intercorrelations based on the number of statistical tests in respective analyses.

### Spatial colocalization analyses

Results of the structural covariance analyses were tested for their topographical colocalization with cholinergic synaptic markers for normative VAChT, M1-mAChR and α4β2-nAChR density maps with Spearman correlation coefficients (*ρ*) and explained variance measures (*R*^2^) were calculated. Assessment of correspondence between cholinergic nucleus region structural covariance maps and cholinergic projection target region topographies was separately quantified for cerebral cortical, subcortical, cerebellar and a whole brain parcellations, as described above for the cholinergic indices. Statistical significance was assessed based on correlations of a structural covariance map with 1000 surrogate null maps for each of the density maps, to account for spatial autocorrelation.

## Results

### Basal forebrain grey matter is linked to cortical and subcortical acetylcholine transporter density beyond direct projection targets

Ch1-4 grey matter was delineated by applying post-mortem cytoarchitectonic maps at a probability threshold of 50% (**Fig 2A**) and average grey matter volume was calculated for voxels within the mask and for each participant. Based on 45 participants from the HCP with independent MRI retest sessions, we found good test-retest reliability for the correlation of Ch1-4 grey matter volume between session 1 and session 2 (Spearman’s *ρ* (44) = .852, *p* < .001, **Fig. 2B**), indicating that Ch1-4 grey matter volume provides a stable structural marker suitable for subsequent analyses. We then conducted a structural covariance analysis with Ch1-4 as seed region, which showed widespread cortical, subcortical and cerebellar regions coupled to Ch1-4 grey matter volume (**Fig. 2C**). Pronounced effects in regions such as amygdala, hippocampus and cingulate cortex confirmed the Ch1-4 structural covariance map as sensitive marker for its known cholinergic projections.

**Figure 2.**
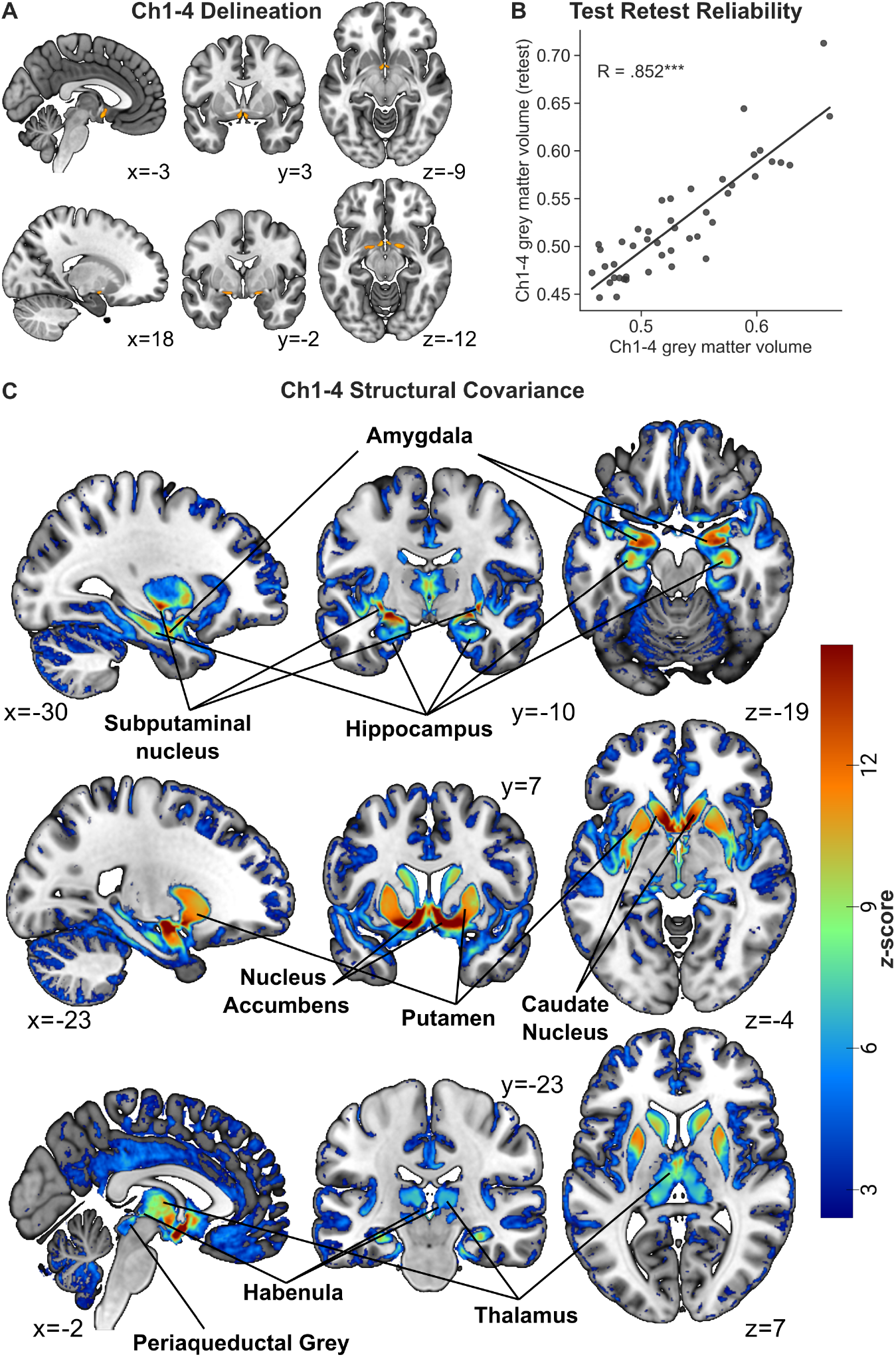
Brain-wide grey matter is coupled with the basal forebrain and exhibits fine-grained regional effects. **A**, The location of the post-mortem cytoarchitectonic maps of the basal forebrain (Ch1-4), as a combined mask for left and right Ch1, Ch2, Ch3 and Ch4 subregions, thresholded at a probability of 50% and presented in orange overlayed on a standard brain template. **B**, Test-retest reliability of Ch1-4 grey matter volume estimates for the 45 participants in the human connectome project cohort which underwent an independent retest magnetic resonance imaging session. Significance is annotated with * for *p* < .05, ** for *p* < .01 and *** for *p* < .001. **C**, Brain-wide grey matter significantly covarying with Ch1-4 grey matter volume, controlled for age, sex and total intracranial volume as additional covariates in the general linear model analysis of the human connectome project cohort. Results are thresholded using a voxel-wise false-discovery rate threshold (*q* < .05) within a 10% grey matter mask and plotted at half maximum for a better contrast.

To examine the validity of Ch1-4 grey matter volume as marker for cholinergic system integrity and its projection targets, we investigated the correspondence between the Ch1-4 structural covariance map and presynaptic VAChT density based on [^18^F]-FEOBV binding from a healthy control group (**Fig. 3A**). In both maps, we saw substantial similarity with the highest values in regions such as striatum, thalamus, periaqueductal grey, habenula, as well as amygdala, hippocampus and cingulate cortex (**Fig. 2C** and **Fig. 3A**). Cortical parcels showed the most pronounced Ch1-4 structural covariance in insular and limbic regions, subcortical parcels in nucleus accumbens, putamen and amygdala, as well as cerebellar parcels in vermis regions (**Fig. 3B-D**). We quantified spatial colocalization of the Ch1-4 structural covariance map with the VAChT density map (**Fig. 3E**), and assessed statistical significance based on the correlation between the Ch1-4 structural covariance map and 1000 surrogate null maps of VAChT density. Significant colocalization was found across the whole brain (voxel-wise: [Spearman’s ρ (51349) = .316, *p_null_* < .001], parcel-wise: [Spearman’s *ρ* (448) =.485, *p_null_* < .001]), as well as across cortical parcels (Spearman’s *ρ* (399) = .397, *p_null_* < .001). Spatial colocalization did not significantly pass control for spatial autocorrelation across subcortical (Spearman’s *ρ* (31) = .411, *p_null_* = .062) and cerebellar (Spearman’s *ρ* (26) = .160, *p_null_* = .638) parcels.

**Figure 3.**
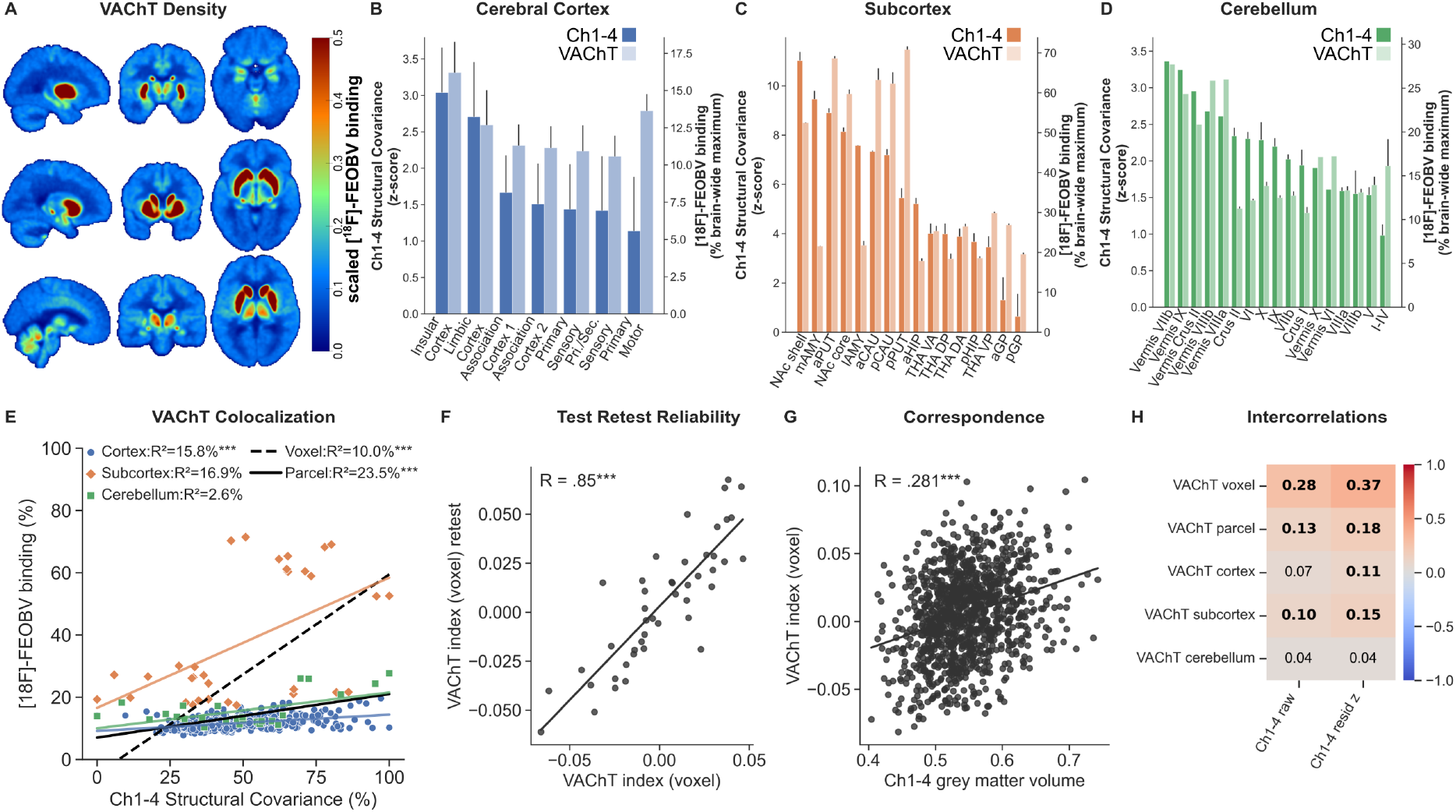
Basal forebrain structural covariance mimics the acetylcholine transporter density map in beyond direct cholinergic projection targets. **A**, Normative radioligand binding of [^18^F]-FEOBV (SUVR) to the presynaptic vesicular acetylcholine transporters (VAChT) in healthy adults, scaled between 0.000001 and 1 within a brain mask, plotted at half maximum for a better contrast and shown in the same brain slices as the Ch1-4 structural covariance map (Fig. 2C). **B-D**, The bar-plots show the distribution within the parcellations for cerebral cortex according to Van-Economo-Koskinos atlas (**B**), subcortical regions according to Tian atlas (**C**) and cerebellar regions according to Diedrichsen atlas (**D**), ordered by their level of Ch1-4 structural covariance effects. Error bars indicate 95% confidence intervals. Note that the y-axes keep the original scales, such as z-scores as in Fig. 2C for structural covariance, and such as scaled binding on to the brain-wide maximum as in Fig. 3A for VAChT density. **E**, Spatial colocalization of the Ch1-4 structural covariance map and normative VAChT density within a 10% grey matter mask assessed by the Spearman correlation of both maps after down-sampling to 3 mm resolution to account for the lower resolution of positron emission tomography data. Significance is controlled for spatial autocorrelation, assessed based on 1000 surrogate null maps and annotated with * for *p* < .05, ** for *p* < .01 and *** for *p* < .001. **F**, Test-retest reliability for a voxel-based VAChT index, which represents the spatial colocalization between VAChT density and a participants’ grey matter volume residuals after correcting for age, sex and intracranial volume within a group-averaged 10% grey matter probability mask. **G**, Correlation between an individuals’ Ch1-4 grey matter and VAChT index for all 1113 participants in the human connectome project cohort. **H**, Correlation between VAChT indices and Ch1-4 measures show descriptively stronger associations with Ch1-4 grey matter when controlled for effects of age, sex and total intracranial volume (tiv_raw_) compared to raw grey matter probability estimates. Significant associations are presented in bold with *p* < .05 controlled for multiple tests using Bonferroni correction.

To extract single participant-level VAChT indices, we determined spatial colocalization between normative VAChT density and an individual’s residual grey matter morphology, after controlling for age, sex and intracranial volume effects. This brain-wide voxel-wise VAChT index had good test-retest reliability between first and replication sessions (Spearman’s *ρ* (44) = .850, *p* < .001; **Fig. 3F**). Individual Ch1-4 grey matter volume and VAChT indices were then correlated in the full HCP sample (Spearman’s *ρ* (1112) = .281, *p* < .001; **Fig. 3G**). The results indicated that an individual’s Ch1-4 volume is linked to the strength of spatial colocalization between that person’s grey matter morphology and normative VAChT density distribution. This link was most pronounced for brain-wide voxel-wise VAChT indices, it was numerically enhanced when age, sex and intracranial volume were residualized from Ch1-4 grey matter volume, and it remained apparent and significant when VAChT indices were restricted to cortical and subcortical parcellations (**Fig. 3H**). We then conducted further analyses to understand whether cholinergic receptor subtypes, such as M1-mAChRs and α4β2-nAChRs, and subregional cholinergic projection targets deriving from Ch1-3 and Ch4 contribute differently to these effects.

### M1-mAChR indices and density highlight subcortical structural covariance coupled to basal forebrain macroanatomy

M1-mAChR receptor density was analyzed based on normative radioligand binding of [^11^C]-LSN3172176 (Naganawa et al., 2021) and depicted the most pronounced binding in the basal ganglia, as well as low levels in brain stem and cerebellum (**Fig. 4A**). Across cortical (**Fig. 4B**), subcortical (**Fig. 4C**) and cerebellar (**Fig. 4D**) regions, nucleus accumbens shell and core, anterior putamen and lateral amygdala expressed the overall highest levels in M1-mAChR receptor density. Spatial colocalization analyses between the Ch1-4 structural covariance map and normative M1-mAChR density was most predominant in subcortical regions (Spearman’s *ρ* (31) = .727, *p_null_* < .001) and significant correspondence was found across the whole-brain for a voxel-wise assessment (Spearman’s *ρ (*51349) = .158, *p_null_* < .001), but not for a parcel-wise assessment (Spearman’s *ρ (*448) = .028, *p_null_* = .748), and not across cortical (Spearman’s *ρ* (399) = .073, *p_null_* = .427) and cerebellar (Spearman’s *ρ* (26) = - .283, *p_null_* = .389) parcels (**Fig. 4E**).

**Figure 4.**
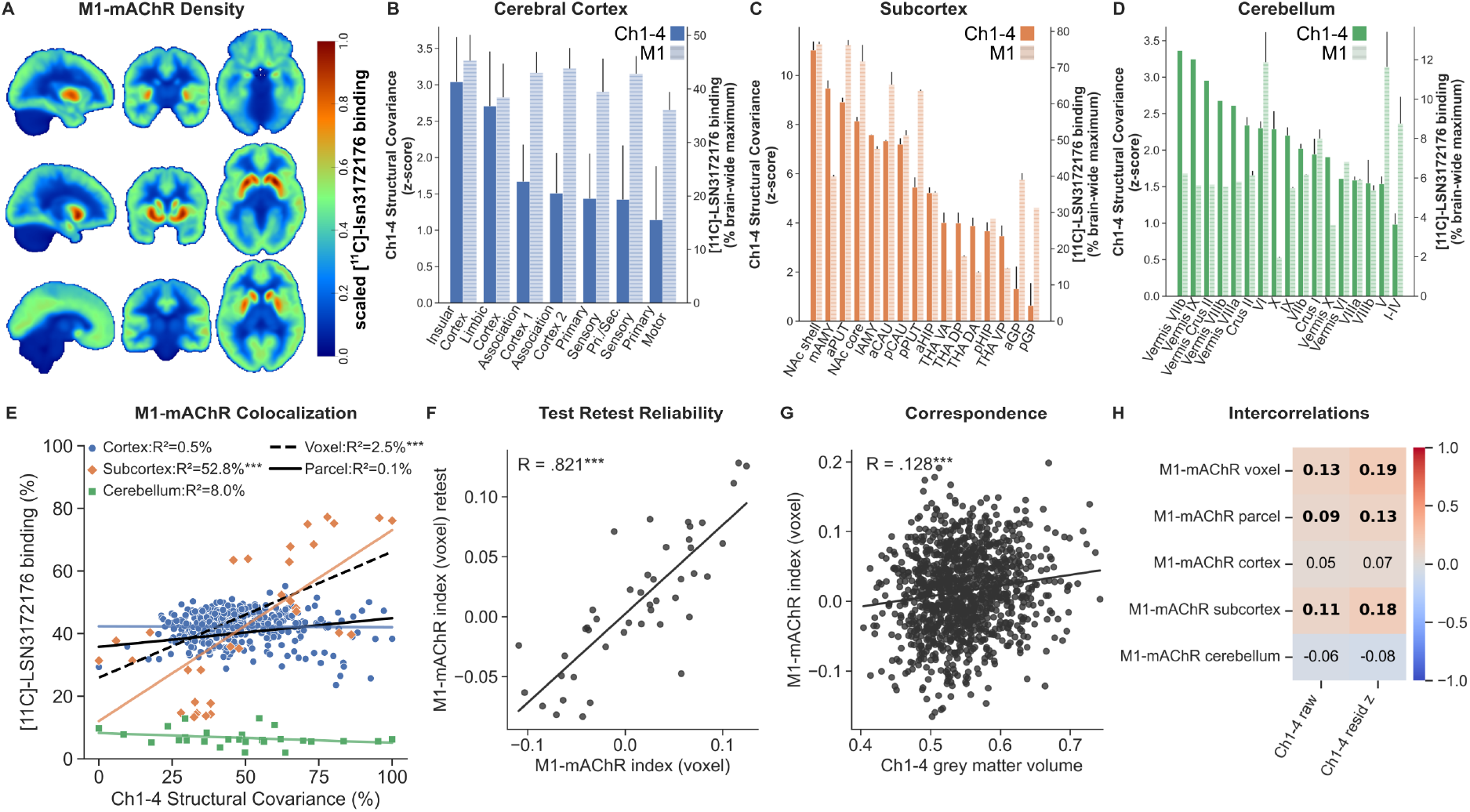
M1-muscarinic acetylcholine receptor density uncovers a subcortical structural covariance gradient most pronounced in nucleus accumbens. **A**, Normative radioligand non- displaceable binding potential (BPND) of the M1-muscarinic acetylcholine receptor (mAChR) ligand [^11^C]-LSN3172176 in healthy adults, scaled between 0.000001 and 1 within a brain mask, plotted at half maximum for a better contrast and shown in the same brain slices as the Ch1-4 structural covariance map density (Fig. 2C). **B-D**, The bar-plots show the distribution within the parcellations for cerebral cortex according to Van-Economo-Koskinos atlas (B), subcortical regions according to Tian atlas (**C**) and cerebellar regions according to Diedrichsen atlas (**D**), ordered by their level of Ch1-4 structural covariance effects. Error bars indicate 95% confidence intervals. Note that the y-axes keep the original scales, such as z-scores as in Fig. 2C for structural covariance, and such as scaled binding on to the brain-wide maximum as in Fig. 4A for M1-mAChR density. **E**, Spatial colocalization of the Ch1-4 structural covariance map and normative M1-mAChR density within a 10% grey matter mask assessed by the Spearman correlation of both maps after down-sampling to 3 mm resolution to account for the lower resolution of positron emission tomography data. Significance is controlled for spatial autocorrelation, assessed based on 1000 surrogate null maps and annotated with * for *p* < .05, ** for *p* < .01 and *** for *p* < .001. **F**, Test-retest reliability for a voxel-based M1-mAChR index, which represents the spatial colocalization between M1-mAChR density and a participants’ grey matter volume residuals after correcting for age, sex and intracranial volume within a group-averaged 10% grey matter probability mask. **G**, Correlation between an individuals’ Ch1-4 grey matter and M1-mAChR index for all 1113 participants in the human connectome project cohort. **H**, Correlation between M1-mAChR indices and Ch1-4 measures show descriptively stronger associations with Ch1-4 grey matter when controlled for effects of age, sex and total intracranial volume (tiv_raw_) compared to raw grey matter probability estimates. Significant associations are presented in bold with *p* < .05 controlled for multiple tests using Bonferroni correction.

Following the same procedure as for VAChT, we found good test-retest reliability for the single-participant voxel-level grey matter-based M1-mAChR index (Spearman’s *ρ* (44) = .821, *p* < .001; **Fig. 4F**). Average Ch1-4 grey matter volume was significantly linked to the participants’ M1-mAChR indices (Spearman’s *ρ* (1112) = .128, *p* < .001; **Fig. 4G**). Residual Ch1-4 grey matter volume, controlled for age, sex and intracranial volume, was associated with brain-wide (voxel- and parcel-wise) and subcortical M1-mAChR indices (**Fig. 4H**).

### α4β2-nicotinic acetylcholine receptor density links basal forebrain grey matter with thalamic and neocortical macrostructure

We next investigated α4β2-nAChR density based on [^18^F]-Flubatine binding (Hillmer et al., 2016), which depicted the pronounced in thalamic, brain stem and cerebellar regions (**Fig. 5A**). Thalamic density dominated the regional assessment of α4β2-nAChR density among all other brain regions and binding was low in those subcortical regions with high M1-mAChR density such as striatal subregions (**Fig. 5B-D)**. Spatial colocalization between α4β2-nAChR density and Ch1-4 structural covariance maps was significant across the whole brain (voxel-wise: [Spearman’s *ρ (*51349) = .205, *p_null_* < .001], parcel-wise: [Spearman’s *ρ (*448) = .225, *p_null_* = .005]), but not across neocortical (Spearman’s *ρ* (399) = .155, *p_null_* =.068), subcortical (Spearman’s *ρ* (31) = -.236, *p_null_* = .323) and cerebellar (Spearman’s *ρ* (26) = .252, *p_null_* = .439) parcellations (**Fig. 5E**).

**Figure 5.**
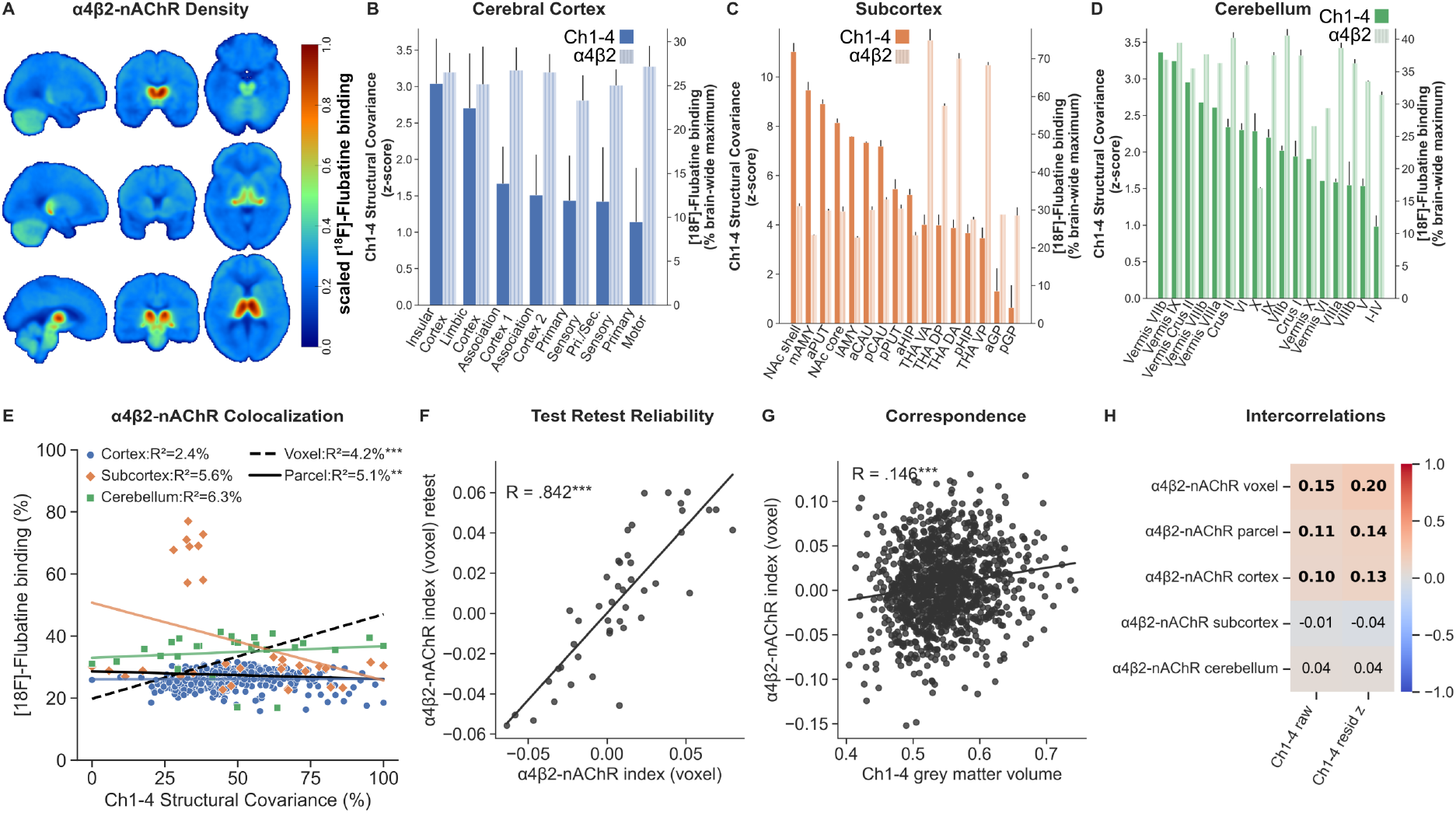
α4β2-nicotinic acetylcholine receptor density is linked to thalamic grey matter and cortical structural covariance gradients of the basal forebrain. **A**, Normative radioligand binding of [^18^F]-Flubatine (VT) to the α4β2-nicotinic acetylcholine receptor (nAChR) in healthy adults, scaled between 0.000001 and 1 within a brain mask, plotted at half maximum for a better contrast and shown in the same brain slices as the Ch1-4 structural covariance map density (Fig. 2C). **B-D**, The bar-plots show the distribution within the parcellations for cerebral cortex according to Van-Economo-Koskinos atlas (B), subcortical regions according to Tian atlas (**C**) and cerebellar regions according to Diedrichsen atlas (**D**), ordered by their level of Ch1-4 structural covariance effects. Error bars indicate 95% confidence intervals. Note that the y-axes keep the original scales, such as z-scores as in Fig. 2C for structural covariance, and such as scaled binding on to the brain-wide maximum as in Fig. 5A for α4β2-nAChR density. **E**, Spatial colocalization of the Ch1-4 structural covariance map and normative α4β2-nAChR density within a 10% grey matter mask assessed by the Spearman correlation of both maps after down-sampling to 3 mm resolution to account for the lower resolution of positron emission tomography data. Significance is controlled for spatial autocorrelation, assessed based on 1000 surrogate null maps and annotated with * for *p* < .05, ** for *p* < .01 and *** for *p* < .001. **F**, Test-retest reliability for a voxel-based α4β2-nAChR index, which represents the spatial colocalization between α4β2-nAChR density and a participants’ grey matter volume residuals after correcting for age, sex and intracranial volume within a group-averaged 10% grey matter probability mask. **G**, Correlation between an individuals’ Ch1-4 grey matter and α4β2-nAChR index for all 1113 participants in the human connectome project cohort. **H**, Correlation between α4β2-nAChR indices and Ch1-4 measures show descriptively stronger associations with Ch1-4 grey matter when controlled for effects of age, sex and total intracranial volume (tiv_raw_) compared to raw grey matter probability estimates. Significant associations are presented in bold with *p* < .05 controlled for multiple tests using Bonferroni correction.

The grey matter-based α4β2-nAChR index showed good test-retest reliability between first and replication sessions (Spearman’s *ρ* (44) = .842, *p* < .001; **Fig. 5F**) and was significantly correlated with an individual’s Ch1-4 grey matter volume (Spearman’s *ρ* (1112) = .146, *p* < .001; **Fig. 5G**). α4β2-nAChR indices were linked to Ch1-4 grey matter volume, when they derived from whole-brain grey matter voxels and parcels, as well as when they were restricted to neocortical regions (**Fig. 5H**).

### Ch1-3 structural covariance uncovers subcortical cholinergic topographies in locations of striatal interneurons and shared fornix connections

The Ch1-3 mask consisted of three cholinergic nuclei, i.e., the medial septum (Ch1), as well as vertical (Ch2) and horizontal limbs (Ch3) of the diagonal band of Broca. We thresholded the cytoarchitectonic Ch1-3 mask at a 50% probability threshold (**Fig. 6A**), and found good test-retest reliability between first and retest sessions (Spearman’s *ρ* (44) = .827, *p* < .001; **Fig. 6B**). Ch1-3 grey matter volume correlated with whole-brain (voxel- and parcel-based) VAChT, M1-mAChR and α4β2-nAChR indices, when controlled for effects of age, sex and intracranial volume (**Fig. 6C**). To determine the specificity for Ch1-3 projection targets, we calculated individual-participant Ch1-3 grey matter volume residuals (Ch1-3_ortho_), by applying statistical orthogonalization of Ch4 grey matter volume. Ch1-3_ortho_ grey matter volume significantly correlated with subcortical VAChT and M1-mAChR indices (**Fig. 6C**), which is consistent with subcortical projection targets such as hippocampus from Ch1 and Ch2. As well, Ch1-3_ortho_ grey matter volume correlated with sex and intracranial volume independent of linear Ch4 effects.

**Figure 6.**
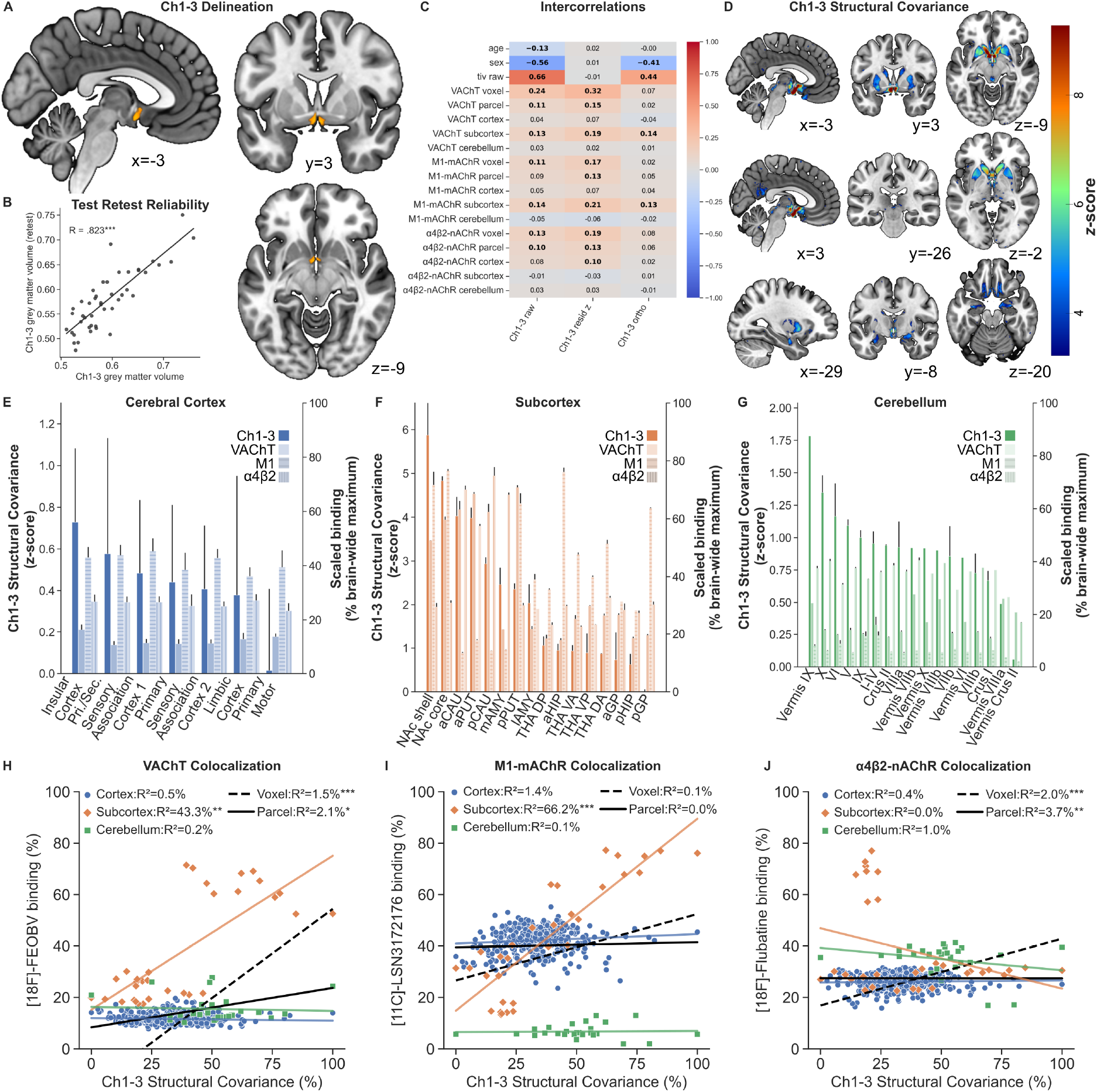
Ch1-3 structural covariance effects colocalize with subcortical acetylcholine transporter and M1-receptor density. **A**, The location of the post-mortem cytoarchitectonic map for medial septum and diagonal bad of Broca (Ch1-3) subregion in the basal forebrain, as a combined mask for left and right Ch1, Ch2 and Ch3 subregions, thresholded at a probability of 50% and presented in orange overlayed on a standard brain template. B, Test-retest reliability of Ch1-3 grey matter volume estimates for the 45 participants in the human connectome project cohort which underwent an independent retest magnetic resonance imaging session. Significance is annotated with * for p < .05, ** for p < .01 and *** for p < .001. C, Intercorrelations between Ch1-3 grey matter measures, VAChT, M1-mAChR and α4β2-nAChR indices, as well as effects of age, sex and total intracranial volume (tivraw). The three columns represent raw grey matter (Ch1-3raw), z-scored grey matter controlled for age, sex and total intracranial volume (Ch1-3resid), and raw grey matter statistically orthogonalized (Ch1-3ortho) from an individuals’ basal nucleus of Meynert/Ch4 grey matter estimate. Significant associations are presented in bold with p < .05 controlled for multiple tests using Bonferroni correction. D, Brain-wide grey matter significantly covarying with orthogonalized Ch1-3 grey matter volume, controlled for age, sex and total intracranial volume as additional covariates in the general linear model analysis of the human connectome project cohort. Results are thresholded using a voxel-wise false-discovery rate threshold (q < .05) within a 10% grey matter mask. E-G, The bar-plots show the distribution within the parcellations for cerebral cortex according to Van-Economo-Koskinos atlas (E), subcortical regions according to Tian atlas (F) and cerebellar regions according to Diedrichsen atlas (G), ordered by their level of Ch1-3 structural covariance effects. Bar plots from left to right indicate: Ch1-3 structural covariance (darker colour), VAChT (lighter colour), M1-mAChR (horizontal lines) and α4β2-nAChR (vertical lines). Error bars indicate 95% confidence intervals. Note that the right y-axis keep the original scales as in (Fig. 3A), (Fig. 4A) and (Fig. 5A) which were all independently scaled to a brain-wide maximum of 100%, and cover up to different values to allow overall regional effect comparisons. H-J, Spatial colocalization of the orthogonalized Ch1-3 structural covariance map and normative VAChT (H), M1-mAChR (I) and α4β2-nAChR (J) density within a 10% grey matter mask assessed by the Spearman correlation of both maps after down-sampling to 3 mm resolution to account for the lower resolution of positron emission tomography data. Significance is controlled for spatial autocorrelation, assessed based on 1000 surrogate null maps and annotated with * for p < .05, ** for p < .01 and *** for p < .001.

The Ch1-3_ortho_ structural covariance map uncovered several further regions, which we previously found to correspond between Ch1-4 structural covariance and VAChT density maps, such as basal ganglia, periaqueductal grey, habenula and amygdala (**Fig. 6D**). Coupled grey matter was, however, also found in areas such as hypothalamus and a cluster extending posterolaterally from the Ch4 mask into claustrum, which overlapped with previous reports on the location of the subputaminal nucleus of Ayala and which had a replicable topography in the Ch1-3_ortho_ structural covariance map of the replication sample (**Supplementary Figure S2**). Regional associations emphasized the most pronounced grey matter coupling with striatal regions, such as nucleus accumbens, caudate nucleus and putamen (**Fig. 6E-G**). Spatial colocalization analyses (**Fig. 6H-J)** largely recapitulated intercorrelations of Ch1-3 grey matter volume with VAChT, M1-mAChR and α4β2-nAChR indices. The Ch1-3_ortho_ structural covariance map had significant spatial colocalization with VAChT density across the whole brain (voxel-wise: [Spearman’s *ρ (*51349) = .123, *p_null_* < .001], parcel-wise: [Spearman’s *ρ (*448) = .146, *p_null_* = .036]) and across subcortical parcels (Spearman’s *ρ* (31) = .658, *p_null_* = .001). M1-mAChR density significantly colocalized with the Ch1-3 structural covariance map across subcortical parcels (Spearman’s *ρ* (31) = .813, *p_null_* < .001), and α4β2-nAChR colocalized across whole-brain grey matter regions (voxel-wise: [Spearman’s *ρ (*51349) = .141, *p_null_* < .001], parcel-wise: [Spearman’s *ρ (*448) = .192, *p_null_ <* .001]). All other spatial colocalization results were non-significant after controlling for spatial autocorrelation (|Spearman’s *ρ*| ≤ .118, *p_null_* ≥ .109).

### Ch4 grey matter is coupled with its corticopetal limbic and insular targets but also with hippocampus and medial thalamus

The basal nucleus of Meynert represents the Ch4 subregion of the basal forebrain and derives corticopetal cholinergic projections to cerebral cortical regions and amygdala. We thresholded the Ch4 mask again at 50% cytoarchitectonic probability (**Fig. 7A**) and found good test-retest reliability between first and retest sessions of the participants in the HCP cohort (Spearman’s *ρ* (44) = .831, *p* < .001; **Fig. 7B**). To delineate effects specific to Ch4, participant’s grey matter volumes were statistically orthogonalized from Ch1-3 effects. Ch4_ortho_ grey matter volume was still associated with whole-brain voxel-wise and cortical VAChT indices (**Fig. 7C**), in line with the predominant cortical cholinergic projection targets of Ch4. As well, Ch4_ortho_ grey matter volume was negatively correlated with age independent of linear Ch1-3 effects.

**Figure 7.**
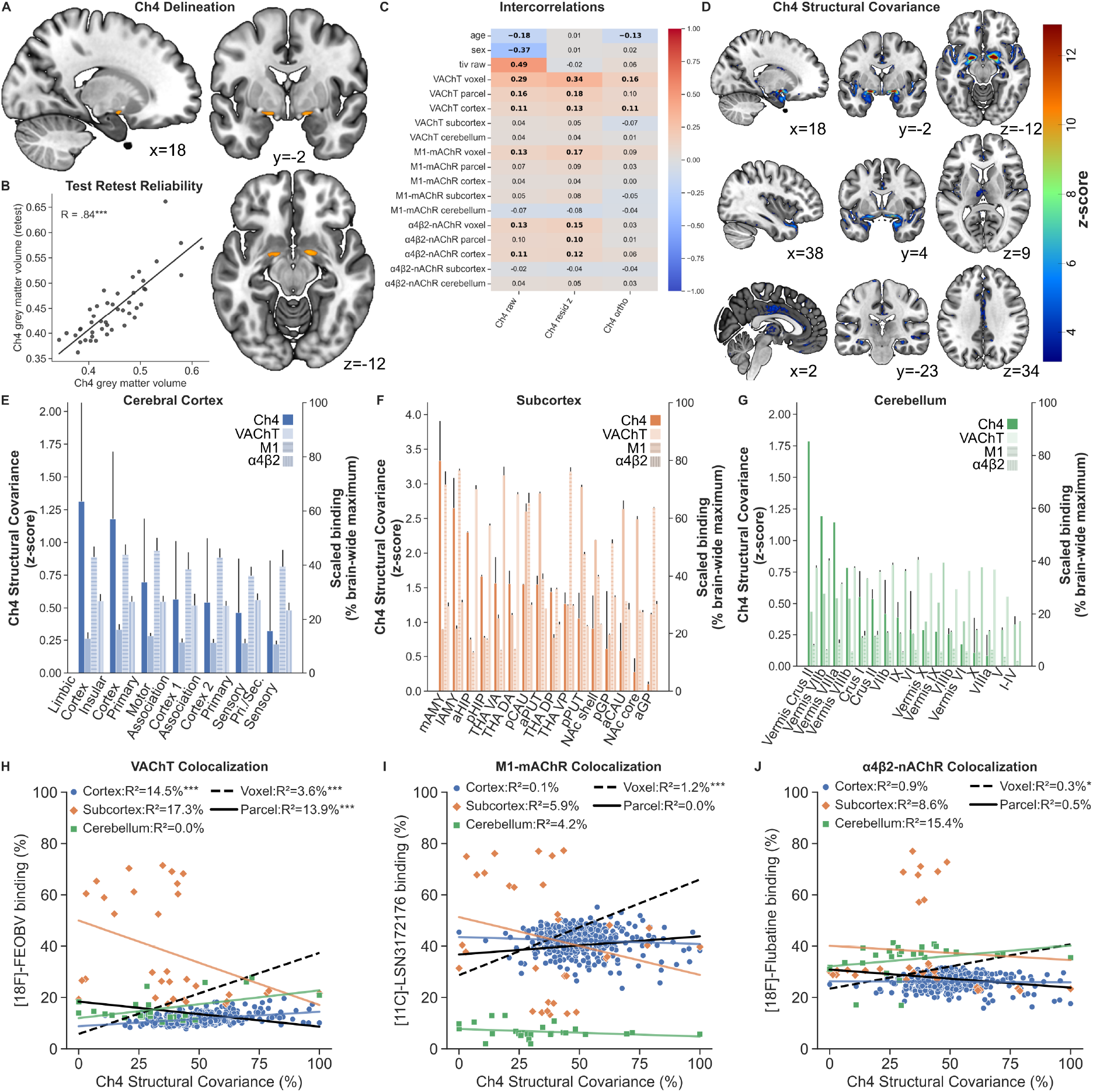
Ch4 structural covariance highlights hippocampal and thalamic effects beyond direct amygdala, insula and cingulate cortex projections. **A**, The location of the post-mortem cytoarchitectonic map for the basal nucleus of Meynert (Ch4) subregion in the basal forebrain, as a combined mask for left and right Ch4 subregions, thresholded at a probability of 50% and presented in orange overlayed on a standard brain template. B, Test-retest reliability of Ch4 grey matter volume estimates for the 45 participants in the human connectome project cohort which underwent an independent retest magnetic resonance imaging session. Significance is annotated with * for p < .05, ** for p < .01 and *** for p < .001. C, Intercorrelations between Ch4 grey matter measures, VAChT, M1-mAChR and α4β2-nAChR indices, as well as effects of age, sex and total intracranial volume (tivraw). The three columns represent raw grey matter (Ch4raw), z-scored grey matter (Ch4resid) controlled for age, sex and total intracranial volume, and raw grey matter statistically orthogonalized from an individuals’ medial septum and diagonal band of Broca/Ch1-3 grey matter estimate (Ch4ortho). Significant associations are presented in bold with p < .05 controlled for multiple tests using Bonferroni correction. D, Brain-wide grey matter significantly covarying with orthogonalized Ch4 grey matter volume, controlled for age, sex and total intracranial volume as additional covariates in the general linear model analysis of the human connectome project cohort. Results are thresholded using a voxel-wise false-discovery rate threshold (q < .05) within a 10% grey matter mask. E-G, The bar-plots show the distribution within the parcellations for cerebral cortex according to Van-Economo-Koskinos atlas (E), subcortical regions according to Tian atlas (F) and cerebellar regions according to Diedrichsen atlas (G), ordered by their level of Ch4 structural covariance effects. Bar plots from left to right indicate: Ch4 structural covariance (darker colour), VAChT (lighter colour), M1-mAChR (horizontal lines) and α4β2-nAChR (vertical lines). Error bars indicate 95% confidence intervals. Note that the right y-axis keep the original scales as in (Fig. 3A), (Fig. 4A) and (Fig. 5A) which were all independently scaled to a brain-wide maximum of 100%, and cover up to different values to allow overall regional effect comparisons. H-J, Spatial colocalization of the orthogonalized Ch4 structural covariance map and normative VAChT (H), M1-mAChR (I) and α4β2-nAChR (J) density within a 10% grey matter mask assessed by the Spearman correlation of both maps after down-sampling to 3 mm resolution to account for the lower resolution of positron emission tomography data. Significance is controlled for spatial autocorrelation, assessed based on 1000 surrogate null maps and annotated with * for p < .05, ** for p < .01 and *** for p < .001.

We conducted a structural covariance analysis with Ch4_ortho_ as seed region, to investigate whether grey matter volume is coupled with its projection targets. Pronounced effects were found in a cluster extending from amygdala to medial parahippocampal gyrus and anterior temporal pole, cingulate cortex, insula, medial thalamus and hippocampus (**Fig. 7D**). Ch1-4 structural covariance effects and regional associations of participants in the HCP exploration cohort were largely replicated in the independent IXI cohort (**Supplementary Figure S3**). To understand whether synaptic cholinergic innervation explains the distribution of the Ch4 structural covariance map, we performed spatial colocalization analyses of the Ch4_ortho_ structural covariance map with VAChT, M1-mAChR and α4β2-nAChR density as described before for the whole basal forebrain and for the Ch1-3 subregion. Within regional comparisons, Ch4_ortho_ structural covariance effects in most regions were considerably dampened by statistical orthogonalization of Ch1-3 grey matter volume (**Fig. 7E-G)**. Still, Ch4_ortho_ structural covariance was strongest in amygdala, limbic and insular cortices, which are assumed to receive the densest synaptic cholinergic projections from Ch4 through medial and lateral corticopetal pathways. The Ch4_ortho_ structural covariance map significantly colocalized with VAChT density across whole-brain grey matter expression (voxel-wise: [Spearman’s *ρ (*51349) = .189, *p_null_* < .001], parcel-wise: [Spearman’s *ρ (*448) = .373, *p_null_ <* .001]) and across cortical parcels (Spearman’s *ρ* (399) = .380, *p_null_* < .001, **Fig. 7H**). For whole-brain voxel-wise colocalization, significant associations of Ch4_ortho_ structural covariance were found for M1-mAChR density (Spearman’s *ρ (*51349) = .108, *p_null_* < .001, **Fig. 7I**) and α4β2-nAChR (Spearman’s *ρ (*51349) = .054, *p_null_* = .023, **Fig. 7J**). All other spatial colocalization analyses were non-significant after controlling for spatial autocorrelation (|Spearman’s *ρ*| ≤ .415, *p_null_* ≥ .088).

## Discussion

In humans, it is assumed that most synaptic cholinergic innervation in neocortex, amygdala and hippocampus derives from Ch1-4 projections (Oda and Nakanishi, 2000; Geula et al., 2025). In these cholinoceptive regions, M1-mAChR and α4β2-nAChR are important molecular targets for synaptic signalling, and VAChT are involved in the reuptake of acetylcholine into presynaptic vesicles. Measuring molecular cholinergic components as synaptic markers for cholinergic target regions in the living brain has previously required invasive and complex procedures. Here, we demonstrated that specific signatures of the cholinergic system are embedded within macroscale gradients of grey matter morphology using commonly acquired anatomical MRI. Specifically, we found good test-retest reliability for Ch1-4, Ch1-3 and Ch4 grey matter volumes, as well as for individual-level indices capturing associations between residual grey matter distribution and normative density patterns of VAChT, M1-mAChR and α4β2-nAChR. Intercorrelations between cholinergic nucleus and target region measures revealed that Ch1-4 grey matter volume captures acetylcholine-specific effects and that cholinergic target region indices vary in their cortical and subcortical contributions. Ch1-4 structural covariance analyses resembled a widespread and fine-grained map indicating various potential cholinergic effects embedded in the brain’s grey matter architecture. By quantifying the spatial colocalization with normative VAChT, M1-mAChR and α4β2-nAChR density maps, we demonstrated that Ch1-4 structural covariance converged in central regions of these synaptic cholinergic topographies, while leveraging the superior spatial resolution of structural MRI. A subcortical grey matter gradient was first indicated by the spatial colocalization of the Ch1-4 structural covariance map with VAChT density and later identified as largely explained by M1-mAChR colocalization. Ch4 grey matter volume was coupled with amygdala, insular and cingulate cortices, confirming regions previously described as having dense cholinergic innervation derived from Ch4 (Selden et al., 1998). Significant Ch1-3 structural covariance effects were found in regions among orbitofrontal cortex, at the intersection of hippocampus and amygdala, as well as other regions that share fornix contributions, such as hypothalamus and retrosplenial cortex. Overall, these findings support the use of grey matter-derived metrics of the cholinergic system in neuroscientific and clinical applications.

### Identification of cholinergic circuit subpopulations

Subregional structural covariance results partially confirmed, but also contradicted previous reports on cholinergic nucleus region projection patterns in certain regions. Spatial colocalization analyses of VAChT, M1-mAChR and α4β2-nAChR density maps provided additional molecular insight into the underlying cholinergic circuits.

While cortical and amygdala effects were expected as direct cholinergic projection targets of Ch4, parahippocampal and cingulate regions also had pronounced structural covariance with Ch4 and were among those previously described limbic and paralimbic regions which back-project to Ch4 in humans (Mesulam, Mufson, Levey, et al., 1983; Mesulam & Geula, 1988). Cholinergic index intercorrelations and spatial colocalization analyses further confirmed that the cortical gradient of synaptic cholinergic innervation aligned with the Ch4 covariance map. This finding supports the idea of Ch4-based grey matter maintenance in cortical synaptic targets or Ch4 maintenance by limbic back-projecting neurons. Ch4 structural covariance effects in medial thalamus were, as well, robust and replicable, although a cholinergic origin driven by Ch5 in pedunculopontine nucleus, Ch6 in laterodorsal tegmental nucleus projections would be more consistent with previous studies in humans (Mesulam et al., 1989) and animals (Mesulam et al., 1983b). A previous study found thalamic cholinergic innervation deriving from Ch4 in humans by tracing retrograde nerve growth factor back-propagation (Heckers et al., 1992), providing potential molecular hints on the origin of grey matter coupling between cholinergic nucleus regions and their synaptic projection targets. Additionally, bilateral hippocampal grey matter coupling was observed despite orthogonalization of Ch1-3 grey matter volumes, and replicated in another MRI cohort. Previous studies have shown that hippocampal innervation from Ch4 passes through entorhinal cortex and alveolar hippocampal input pathway in humans, although it has been suggested that this portion is minor (Ransmayr et al., 1989). Given that the age range of our participants only reached up to 40 years in the exploration sample, the results were unlikely to be based on neurodegenerative pathologies, for which a molecular spread of pathological proteins from Ch4 to entorhinal cortex and hippocampus has been found and suggested to precede hippocampal atrophy in Alzheimer’s disease (Cantero et al., 2016; Schmitz et al., 2018; Fernández-Cabello et al., 2020). These coupling patterns are important to consider from a clinical perspective, because cholinesterase inhibitors represent a primary molecular target of anti-dementia treatments for these conditions. Therefore, the results prompt further studies to determine mechanistic links underlying grey matter coupling of Ch4 with hippocampus and thalamus. Further studies on clinical samples are needed to better understand whether Ch4-hippocampal grey matter coupling diminishes or enhances in certain neurodegenerative pathologies, which may emphasize potential cholinergic denervation. Diffusion-based MRI may provide a methodology to investigate whether tract-based metrics on white matter pathways connecting Ch4 to hippocampus and thalamus are also related to Ch4 grey matter volume, or if the associations are best explained by the large proportion of cholinergic neurons in Ch4 compared to Ch1-3 subregions.

One of the regions, which had a robust and replicable structural covariance association across subregions and across MRI cohorts, was found posterolaterally outside the Ch4 mask. It was located at the lowest portion of putamen and extended into a fine line of claustrum. Based on its location, we assumed that it might be best identified as subputaminal nucleus of Ayala, which has been suggested to be a poorly understood, yet human-enriched cholinergic cell group relevant to neurodegenerative pathologies such as Alzheimer’s disease and primary progressive aphasia (Sĭmić et al., 1999; Boban et al., 2006; Teipel et al., 2014). Localization of subputaminal nucleus in humans follows anatomical landmarks, such as starting in coronal brain section from an anterior border at the anterior commissure and a posterior border at the mammillary bodies, where the posterior portion of the subputaminal nucleus is located medial to the putamen and lateral to the anterior commissure (Ayala, 1915; Hamodat et al., 2019). While the structural covariance effects overlapped with this location at its posterior border, about half of the effect also spanned further posterior from the mammillary bodies, which is outside the suggested anatomical landmark (Ayala, 1915; Hamodat et al., 2019). Therefore, the analyses are inconclusive regarding a specific identification of cholinergic cell groups belonging to subputaminal nucleus. While MRI-based localization benefits from submillimeter resolution, it requires more explicit molecular evidence for the confirmation of a cholinergic nucleus region at subputaminal location. PET-based colocalization analyses on the radioligands investigated in this study provide unique insight into specific molecular targets of acetylcholine *in vivo*. However, they are insufficient for confirming subputaminal nucleus of Ayala given its small size estimated below 2 mm^3^ (Grinberg and Heinsen, 2007). On the over hand, in posterior locations subputaminal nucleus has been described to have multiple heterogenous cluster extensions (Ayala, 1915), which may increase MRI-based detectability. More generally, to evaluate such small brain structures and validate the delineation of cholinergic circuits for each participant, post-mortem immunohistochemistry using choline acetyltransferase staining, or targeting synaptic components such as VAChT or acetylcholine esterase, may be better validation approaches when used alongside *in vivo* MRI and single-participant structural covariance analyses.

For Ch1-3 structural covariance, we expected to find pronounced structural coupling with hippocampus since it receives cholinergic projections from medial septum and vertical limb of diagonal band of Broca (Mesulam, Mufson, Levey, et al., 1983; Mesulam & Geula, 1988). However, the strongest effects were found in nucleus accumbens, putamen and caudate nucleus. Coupling in these regions persisted even after control for Ch4 grey matter volume, age, sex and intracranial volume, indicating the robustness of their associations. Additionally, these regions were found to have some of the highest densities of VAChT and M1-mAChR. Cholinergic synapses in these regions are generally believed to be independent of projections from Ch1-8 nuclei and instead solely based on striatal cholinergic interneurons (Oda and Nakanishi, 2000; Geula et al., 2025). Furthermore, M1-mAChRs play a crucial role for striatal interneurons and their synaptic regulation of mesolimbic dopaminergic projections. Because Ch1-3 covariance and M1-mAChR exhibited substantial spatial colocalization across subcortical parcels and in striatal regions, it is reasonable to assume that striatal cholinergic interneurons and their trophic similarity profiles were captured by Ch1-3 covariance. However, more conclusive evidence on the location and effects of these interneurons may require either the successful development of methodologies for localizing cholinergic neurons using only MRI (Wang et al., 2022) or post-mortem histology of individuals who previously underwent MRI scanning. Assessing the effects of striatal cholinergic interneurons in humans could provide significant translational value for understanding the mechanisms of schizophrenia, as these neurons modulate mesolimbic dopaminergic projections and a recently approved muscarinic antipsychotic drug preferentially targets M1- and M4-mAChR receptors (Kaul et al., 2024a, 2024b).

Previous studies applied similar Ch1-4 delineation procedures to explore cholinergic system involvement in neuropsychiatric conditions such as psychosis (Avram et al., 2021; Schulz et al., 2023), Alzheimer’s disease (Schmitz and Spreng, 2016; Fernández-Cabello et al., 2020; Mieling et al., 2023), Parkinson’s disease (Kamalkhani and Zarei, 2022; Slater et al., 2024; Eisenstein et al., 2026), dementia with Lewy bodies (Schumacher et al., 2021; Rau et al., 2025), mild cognitive impairment (Muth et al., 2010; Teipel et al., 2011; Grothe et al., 2014, 2016; Cantero et al., 2016; Schumacher et al., 2021; Shafiee et al., 2024) and subjective cognitive decline (Scheef et al., 2019). Due to the low molecular specificity of MRI, the need to validate the cholinergic basis of Ch1-4 macrostructural measures has been emphasized (Selden et al., 1998). The current results generally support Ch1-4 and subregional grey matter volume as sensitive markers for cholinergic system status, since their interindividual variation corresponded with brain-wide cholinergic innervation patterns. These patterns captured important direct cholinergic projection targets, such as cingulate cortex and amygdala. However, we caution against interpreting that subregional grey matter volume in Ch1-3 and Ch4 as specific to either projection targets.

Previous studies have found alterations in Ch1-4 or its subregions in neuropsychiatric conditions whose symptoms are more closely associated with other cholinergic subpopulations. These alterations may be explained by shared developmental influences, vulnerability, or interconnections. As discussed above, diffusion MRI studies are needed to understand whether the finding of thalamic covariance with Ch4 grey matter volume reflects their direct connections in humans or whether grey matter coupling is driven by the shared variance of Ch4 with Ch5/Ch6 integrity and their thalamic cholinergic projections. Ch5 and Ch6 pathologies may be more likely affected by Parkinson’s disease and contribute to motor dysfunctions and REM sleep disorder (Sharma et al., 2025) as well as regulation of mesolimbic dopamine release (Dautan et al., 2016). This is in contrast to the cognitive deficits typically associated with Ch1-4 degeneration. Projections from the medial habenula to the interpeduncular nucleus, originating from Ch7, have been shown to play a crucial role in regulating addiction-related withdrawal symptoms (Salas et al., 2009). However, direct grey matter imaging of Ch5, Ch6, and Ch8 in the parabigeminal nucleus is challenging due to the low proportion of grey matter compared to white matter in the brainstem. Further advances in MRI resolution could enable in vivo mapping of magnocellular cholinergic neurons (Wang et al., 2022) and, once surpassing spatial resolution limitations, help to disentangle selective subregion-specific grey matter coupling patterns.

### Complementary synaptic cholinergic grey matter indices

To increase imaging-based insights and enable clinical applications regarding the grey matter status of the cholinergic system, we introduced receptor and transporter indices for a more complementary assessment of cholinergic nucleus and target regions. We found VAChT, M1-mAChR and α4β2-nAChR indices to have good test-retest reliability, as well as significant intercorrelations with the more established Ch1-4 grey matter measure. These indices showed correlation patterns with Ch1-4, Ch1-3, and Ch4, which largely recapitulated the results of spatial colocalization analyses, which supports the indices’ external validity and the robustness of the findings. Previous studies have proposed cholinergic indices for blood and cerebrospinal fluid samples (Karami et al., 2019, 2021) and for pharmacological electroencephalography challenges (Simpraga et al., 2017, 2018), to develop surrogate measures for pharmacological responding to cholinergic drugs. Single-participant measures, such as subregional Ch1-3 and Ch4 grey matter volume, as well as average tract-based integrity metrics of corticopetal and fornix projections have been used as cholinergic indices, though few studies have labelled them as such (Qiu et al., 2023). Synaptic target region indices for VAChT, M1-mAChR, and α4β2-nAChR can be quantified from standard structural MRI scans. Therefore, they impose no additional time or costs for participant recruitment.

VAChT indices may be applied as surrogate markers for overall cholinergic innervation as inferred from their presynaptic expression and may support pharmacological responder analyses on cholinesterase inhibitor interventions. VAChT indices were generally correlated with Ch1-4 grey matter volume, although the level of shared variance suggests that both indices capture independent variance as well. It is tempting to speculate whether Ch1-4 or VAChT indices would provide earlier sensitivity as cholinergic biomarkers for certain neurodegenerative pathologies.

With recently approved muscarinic antipsychotics having high affinity to M1-mAChR (Kaul et al., 2024a, 2024b), the here proposed M1-mAChR indices could help inform pharmacotherapeutic decisions between conventional dopaminergic and newer cholinergic antipsychotics. Compared to previous mAChR-EEG indices that used scopolamine administration as pharmacological model for cognitive impairments (Simpraga et al., 2017), more selective M1-mAChR ligands such as biperiden (Blokland, 2022) and the M1-mAChR indices introduced in the current study could better elucidate the neuroscientific mechanisms and receptor-specific effects underlying acetylcholine-vulnerable cognitive impairment.

Previous nicotinic indices satisfied the criterion of receptor-specific effects using mecamylamine (Simpraga et al., 2018), a selective α4β2-nAChR antagonist. Although electroencephalography and structural MRI indices may diverge as short-term state and long-term trait measures, the indices proposed here may easily be extended to quantify cholinergic contributions in functional MRI maps of individual participants. In neuropsychiatric conditions such as nicotine dependence and substance use disorders, where α4β2-nAChR modulation mesolimbic dopamine release plays a role, α4β2-nAChR may help differentiate which individuals are more likely to respond to anti-addictive drugs, such as mecamylamine.

Importantly, the current study did not include neuropsychiatric patients, such that clinical implications generally remain speculative. Building on this procedure, future studies with clinical samples or pharmacological challenges could complement the assessment of Ch1-4 nucleus regions with indices of cholinergic target regions. These studies could improve our understanding of cholinergic involvement in cognitive functions, such as memory-related selective attention allocation (Weuthen et al., 2025), motor sequence learning (Voegtle et al., 2024), midcingulo-insular salience network control (Chakraborty et al., 2024b), and overall neurodevelopmental trajectories (Lotter et al., 2024). Given their high retest reliability and complementary associations, such as a pronounced anticorrelation of M1-mAChR and α4β2-nAChR across subcortical regions, the current results encourage the application of cholinergic indices for VAChT, M1-mAChR, and α4β2-nAChR.

### Outlook and Conclusions

M1-mAChR and α4β2-nAChR density explained complementary contributions in Ch1-4 grey matter coupling maps, in contrast to VAChT topography as overall presynaptic cholinergic marker. While neuronal cell diversity in Ch1-4 exhibits substantial variation in neocortical projection patterns (Chakraborty et al., 2024a), other neuromodulatory neurotransmitter systems may have as well contributed to the observed findings, given that they partially target similar synapses and consequently have overlapping integrity estimates (Wearn et al., 2024). To understand the specificity of these findings, it is crucial to investigate the convergences and distinctions of cholinergic system indices with markers of other neuromodulator systems, such as noradrenaline, dopamine, serotonin, and histamine, in their nucleus and target regions. Additionally, addressing neurotransmitter interactions of multiple molecular targets without radiation exposure will likely require non-invasive imaging procedures such as MRI. Respective studies will be essential for the understanding of why pharmacological targeting of above neuromodulators can alleviate similar neuropsychiatric symptoms in conditions such as major depressive disorder. As further normative density maps become available for α7-nAChR, M2-mAChR and M4-mAChR, we anticipate a more comprehensive understanding of cholinergic receptor architecture in humans and look forward to studies investigating how neurotransmitter interactions shape structure and function of the brain in health and disease.

## Data availability

Both used MRI datasets, HCP (https://www.humanconnectome.org/study/hcp-young-adult/document/1200-subjects-data-release) and IXI (https://brain-development.org/ixi-dataset/), are publicly available. Untresholded group-level structural covariance maps derived from the current results have been uploaded on a public repository (https://cloud.uni-jena.de/s/sDzSxpJeRmSD8HP). Analysis code will be available online at the time of publication.

## Acknowledgements

This work was supported by the German Center for Mental Health (DZPG) funded by Bundesministerium für Forschung, Technologie und Raumfahrt (BMFTR) FKZ 01EE2601J (A.W.) and FKZ 01EE2505A (M.W.), and by a Clinician Scientist Grant to D.L.B (IZKF, CSP032). We thank all participants and researchers who have contributed to the recruitment and recording of the HCP, IXI datasets, as well as the neuroimaging community contributing to the public availability of MRI data processing tools and cholinergic system maps.

## Author contributions

A.W., S.C., B.B. and M.W. designed research, A.W. and S.C. performed research, A.W. analyzed data, A.W. wrote the first draft of the paper, A.W., S.C., D.L.B., M.L., B.B. and M.W. edited the paper.

---

**Supplementary Figure S1.**
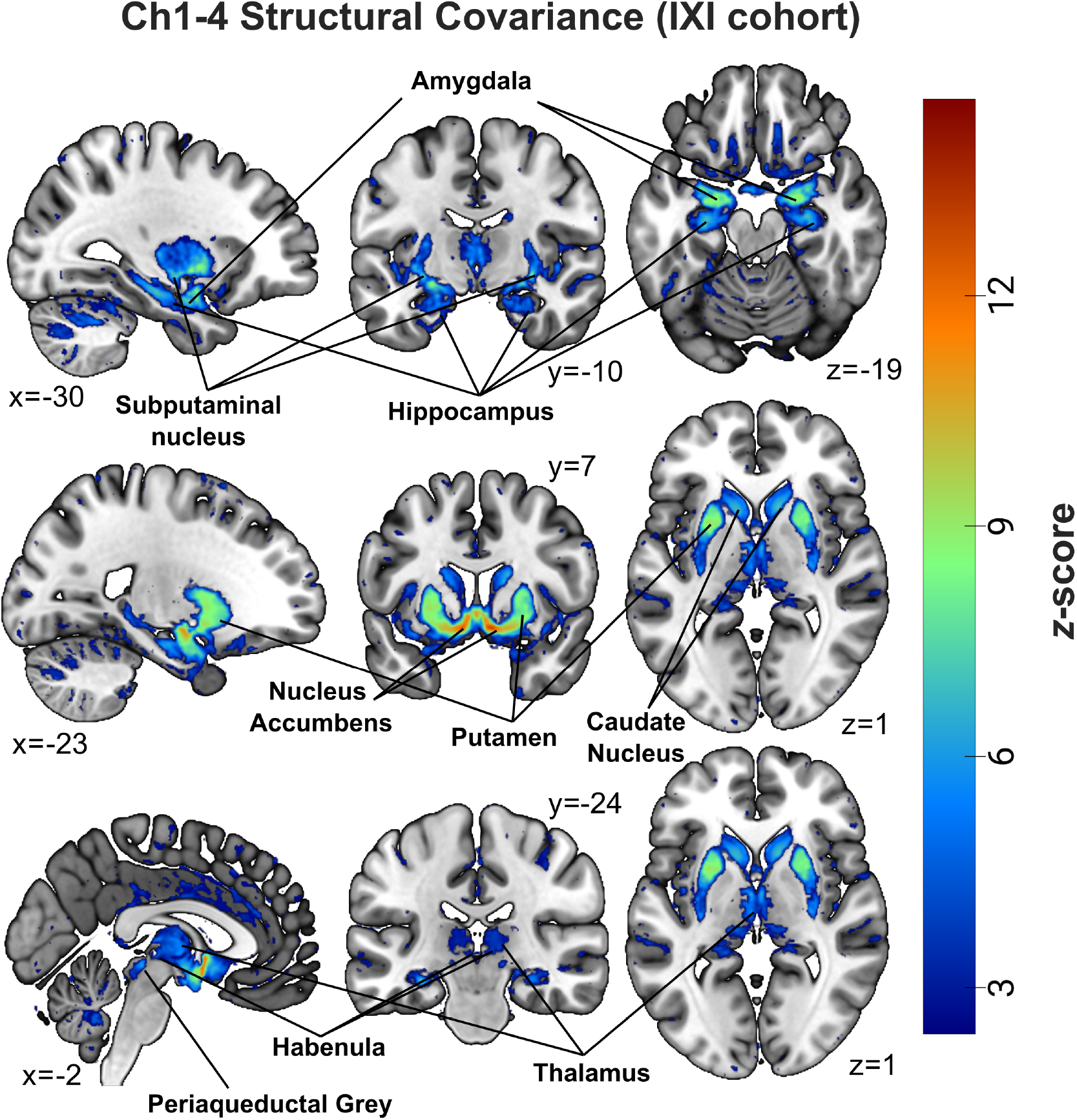
Replication of the Ch1-4 structural covariance analysis in an independent cohort. Brain-wide grey matter covarying with Ch1-4 grey matter volume, controlled for age, sex, total intracranial volume and site as additional covariates in the general linear model analysis. For a better comparability with the analyses from the HCP cohort, results are plotted at the same brain coordinates and visualized using the same color map range.

**Supplementary Figure S2.**
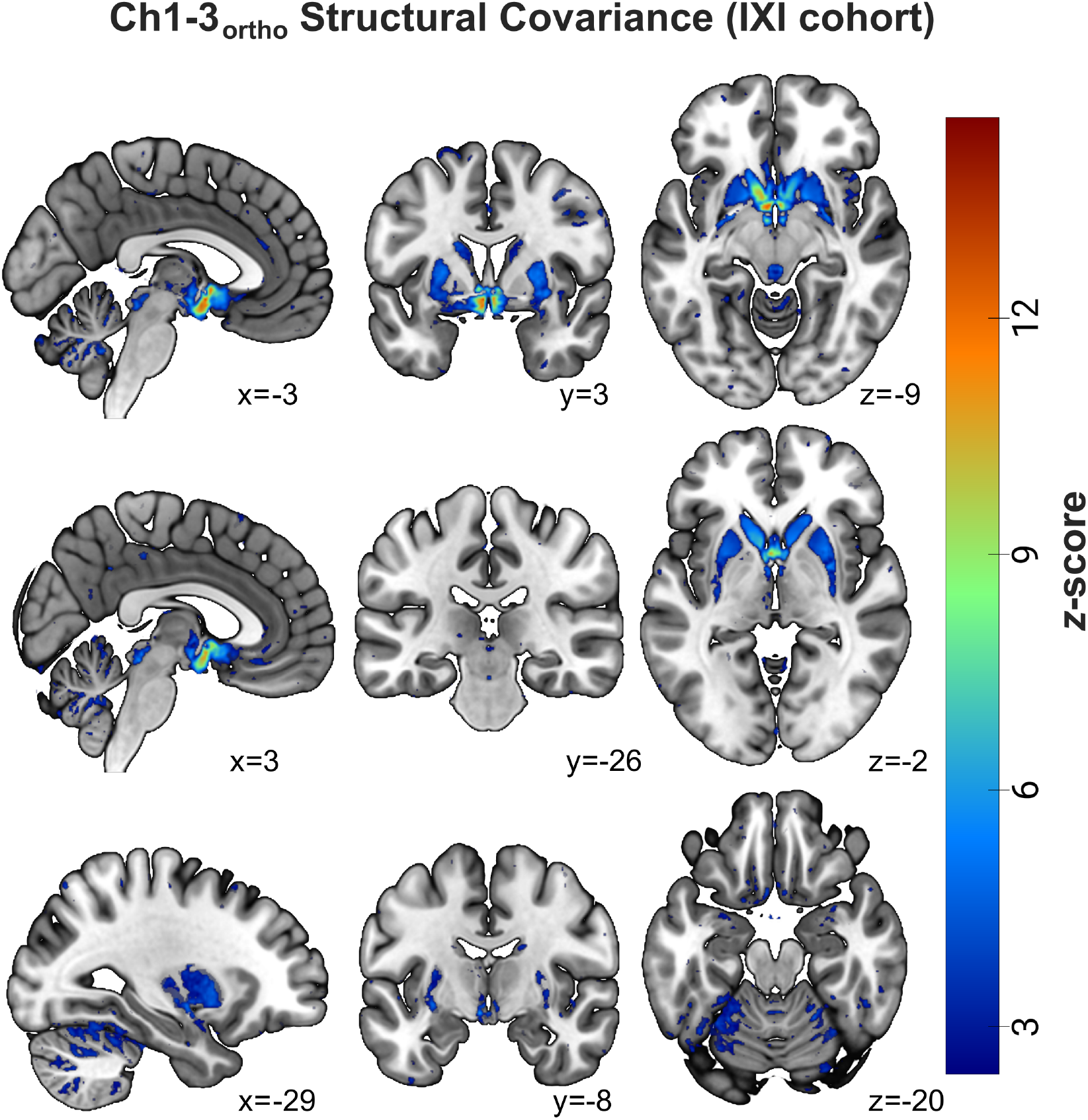
Replication of the Ch1-3_ortho_ structural covariance analysis in an independent cohort. Brain-wide grey matter covarying with orthogonalized Ch1-3 grey matter volume, controlled for age, sex, total intracranial volume and site as additional covariates in the general linear model analysis. For a better comparability with the analyses from the HCP cohort, results are plotted at the same brain coordinates and visualized using the same color map range.

**Supplementary Figure S3.**
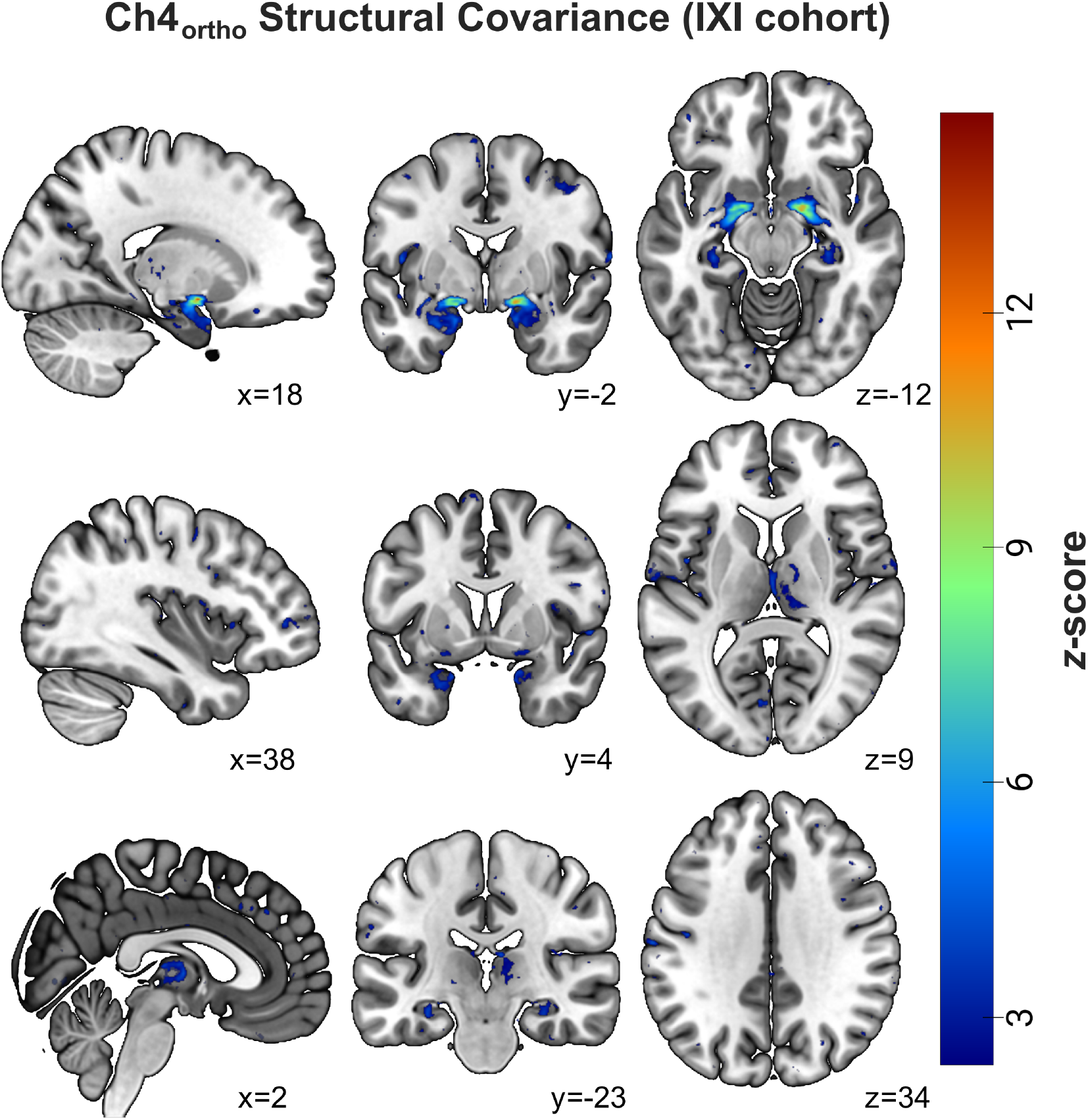
Replication of the Ch1-3ortho structural covariance analysis in an independent cohort. Brain-wide grey matter significantly covarying with orthogonalized Ch4 grey matter volume, controlled for age, sex, total intracranial volume and site as additional covariates in the general linear model analysis. For a better comparability with the analyses from the HCP cohort, results are plotted at the same brain coordinates and visualized using the same color map range.

**Supplementary Figure S4.**
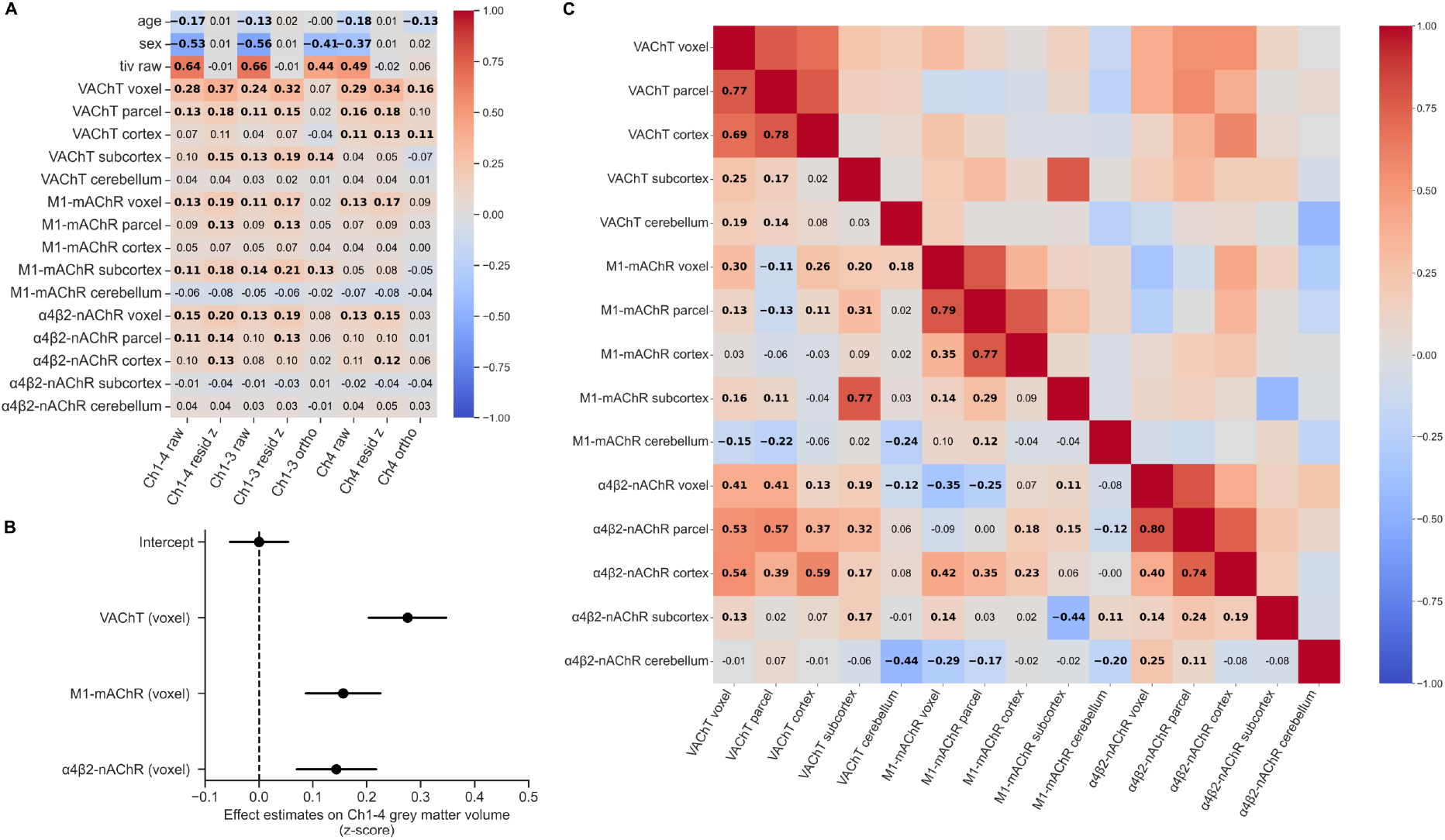
Intercorrelations between cholinergic nucleus and target region indices. **A**, Spearman correlation coefficients between Ch1-4 and subregional nucleus grey matter volume and target region indices of VAChT, M1-mAChR and α4β2-nAChR, as well as age sex and total intracranial volume (tiv_raw_), and controlled for multiple comparison using Bonferroni correction across the table. **B**, General linear model results predicting Ch1-4 grey matter volume based on VAChT, M1-mAChR and α4β2-nAChR indices. **C**, Spearman correlation coefficients for all target region cholinergic indices.

## References

1. Aghourian M, Legault-Denis C, Soucy J-P, Rosa-Neto P, Gauthier S, Kostikov A, Gravel P, Bédard M-A (2017) Quantification of brain cholinergic denervation in Alzheimer’s disease using PET imaging with [18F]-FEOBV. Mol Psychiatry 22:1531–1538.

2. Amunts K, Mohlberg H, Bludau S, Zilles K (2020) Julich-Brain: A 3D probabilistic atlas of the human brain’s cytoarchitecture. Science 369:988–992.

3. Avram M, Grothe MJ, Meinhold L, Leucht C, Leucht S, Borgwardt S, Brandl F, Sorg C (2021) Lower cholinergic basal forebrain volumes link with cognitive difficulties in schizophrenia. Neuropsychopharmacology 46:2320–2329.

4. Ayala G (1915) A hitherto undifferentiated nucleus in the forebrain (nucleus subputaminalis). Brain 37:433–448.

5. Blokland A (2022) Cholinergic models of memory impairment in animals and man: scopolamine vs. biperiden. Behav Pharmacol 33:231–237.

6. Boban M, Kostovic I, Simic G (2006) Nucleus subputaminalis: neglected part of the basal nucleus of Meynert. Brain 129:E42.

7. Bruel-Jungerman E, Lucassen PJ, Francis F (2011) Cholinergic influences on cortical development and adult neurogenesis. Behav Brain Res 221:379–388.

8. Burt JB, Helmer M, Shinn M, Anticevic A, Murray JD (2020) Generative modeling of brain maps with spatial autocorrelation. NeuroImage 220:117038.

9. Cantero JL, Zaborszky L, Atienza M (2016) Volume Loss of the Nucleus Basalis of Meynert is Associated with Atrophy of Innervated Regions in Mild Cognitive Impairment. Cereb Cortex 27:3881–3889.

10. Chakraborty S, Haast RAM, Onuska KM, Kanel P, Prado MAM, Prado VF, Khan AR, Schmitz TW (2024a) Multimodal gradients of basal forebrain connectivity across the neocortex. Nat Commun 15:8990.

11. Chakraborty S, Lee SK, Arnold SM, Haast RAM, Khan AR, Schmitz TW (2024b) Focal acetylcholinergic modulation of the human midcingulo-insular network during attention: Meta-analytic neuroimaging and behavioral evidence. J Neurochem 168:397–413.

12. Ciric R, Thompson WH, Lorenz R, Goncalves M, MacNicol EE, Markiewicz CJ, Halchenko YO, Ghosh SS, Gorgolewski KJ, Poldrack RA, Esteban O (2022) TemplateFlow: FAIR-sharing of multi-scale, multi-species brain models. Nat Methods 19:1568–1571.

13. Crowley SJ, Kanel P, Roytman S, Bohnen NI, Hampstead BM (2024) Basal forebrain integrity, cholinergic innervation and cognition in idiopathic Parkinson’s disease. Brain 147:1799–1808.

14. Cuello AC, Do Carmo S (2025) The dependence of basal forebrain cholinergic neurons on NGF: The case in Alzheimer pathology. In: Cholinergic Involvement in Neurodegenerative Diseases (Cuello AC, ed), pp 95–122 Handbook of Clinical Neurology. Elsevier.

15. Dale HH (1914) The action of certain esters and ethers of choline, and their relation to muscarine. J Pharmacol Exp Ther 6:147–190.

16. Dautan D, Souza AS, Huerta-Ocampo I, Valencia M, Assous M, Witten IB, Deisseroth K, Tepper JM, Bolam JP, Gerdjikov TV, Mena-Segovia J (2016) Segregated cholinergic transmission modulates dopamine neurons integrated in distinct functional circuits. Nat Neurosci 19:1025–1033.

17. Diedrichsen J, Balsters JH, Flavell J, Cussans E, Ramnani N (2009) A probabilistic MR atlas of the human cerebellum. NeuroImage 46:39–46.

18. Eisenstein T, Groenewald K, van Hillegondsberg L, Al Hajraf F, Zerenner T, Lawton MA, Ben-Shlomo Y, Griffanti L, Hu MT, Klein JC (2026) Cholinergic degeneration in prodromal and early Parkinson’s disease: a link to present and future disease states. Brain 149:226–237.

19. Fernández-Cabello S, Kronbichler M, Van Dijk KRA, Goodman JA, Spreng RN, Schmitz TW, on behalf of the Alzheimer’s Disease Neuroimaging Initiative (2020) Basal forebrain volume reliably predicts the cortical spread of Alzheimer’s degeneration. Brain 143:993–1009.

20. Gaser C, Dahnke R, Thompson PM, Kurth F, Luders E, the Alzheimer’s Disease Neuroimaging Initiative (2024) CAT: a computational anatomy toolbox for the analysis of structural MRI data. GigaScience 13.

21. Geula C, Ayala I, Gefen T, Mesulam M-M (2025) Organization of the basal forebrain cholinergic system in the human brain. In: Cholinergic Involvement in Neurodegenerative Diseases (Cuello AC, ed), pp 11–21 Handbook of Clinical Neurology. Elsevier.

22. Gilmore AD, Buser NJ, Hanson JL (2021) Variations in structural MRI quality significantly impact commonly used measures of brain anatomy. Brain Inform 8:7.

23. Grinberg LT, Heinsen H (2007) Computer-assisted 3D reconstruction of the human basal forebrain complex. Dement Neuropsychol 1:140–146.

24. Grothe MJ, Ewers M, Krause B, Heinsen H, Teipel SJ (2014) Basal forebrain atrophy and cortical amyloid deposition in nondemented elderly subjects. Alzheimers Dement 10:344–353.

25. Grothe MJ, Heinsen H, Amaro E Jr, Grinberg LT, Teipel SJ, for the Alzheimer’s Disease Neuroimaging Initiative (2016) Cognitive Correlates of Basal Forebrain Atrophy and Associated Cortical Hypometabolism in Mild Cognitive Impairment. Cereb Cortex 26:2411–2426.

26. Hamodat H, Fisk JD, Darvesh S (2019) Cholinergic Neurons in Nucleus Subputaminalis in Primary Progressive Aphasia. Can J Neurol Sci 46:174–183.

27. Hansen JY, Cauzzo S, Singh K, García-Gomar MG, Shine JM, Bianciardi M, Misic B (2024) Integrating brainstem and cortical functional architectures. Nat Neurosci 27:2500–2511.

28. Heckers S, Geula C, Mesulam M-M (1992) Cholinergic innervation of the human thalamus: Dual origin and differential nuclear distribution. J Comp Neurol 325:68–82.

29. Hillmer AT, Esterlis I, Gallezot JD, Bois F, Zheng MQ, Nabulsi N, Lin SF, Papke RL, Huang Y, Sabri O, Carson RE, Cosgrove KP (2016) Imaging of cerebral α4β2* nicotinic acetylcholine receptors with (−)-[18F]Flubatine PET: Implementation of bolus plus constant infusion and sensitivity to acetylcholine in human brain. NeuroImage 141:71–80.

30. Hohmann CF (2003) A morphogenetic role for acetylcholine in mouse cerebral neocortex. Neurosci Biobehav Rev 27:351–363.

31. Kamalkhani N, Zarei M (2022) Distinct atrophy of septal nuclei in Parkinson’s disease. Clin Park Relat Disord 7:100171.

32. Karami A, Darreh-Shori T, Schultzberg M, Eriksdotter M (2021) CSF and Plasma Cholinergic Markers in Patients With Cognitive Impairment. Front Aging Neurosci 13:704583.

33. Karami A, Eriksdotter M, Kadir A, Almkvist O, Nordberg A, Darreh-Shori T (2019) CSF Cholinergic Index, a New Biomeasure of Treatment Effect in Patients With Alzheimer’s Disease. Front Mol Neurosci 12:239.

34. Kaul I, Sawchak S, Correll CU, Kakar R, Breier A, Zhu H, Miller AC, Paul SM, Brannan SK (2024a) Efficacy and safety of the muscarinic receptor agonist KarXT (xanomeline–trospium) in schizophrenia (EMERGENT-2) in the USA: results from a randomised, double-blind, placebo-controlled, flexible-dose phase 3 trial. The Lancet 403:160–170.

35. Kaul I, Sawchak S, Walling DP, Tamminga CA, Breier A, Zhu H, Miller AC, Paul SM, Brannan SK (2024b) Efficacy and Safety of Xanomeline-Trospium Chloride in Schizophrenia: A Randomized Clinical Trial. JAMA Psychiatry 81:749–756.

36. Kilimann I, Grothe M, Heinsen H, Alho EJL, Grinberg L, Amaro Jr. E, dos Santos GAB, da Silva RE, Mitchell AJ, Frisoni GB, Bokde ALW, Fellgiebel A, Filippi M, Hampel H, Klöppel S, Teipel SJ (2014) Subregional Basal Forebrain Atrophy in Alzheimer’s Disease: A Multicenter Study. J Alzheimer’s Dis 40:687–700.

37. Loewi O (1922) Über humorale Übertragbarkeit der Herznervenwirkung. Pflüg Arch Für Gesamte Physiol Menschen Tiere 193:201–213.

38. Lotter LD et al. (2024) Regional patterns of human cortex development correlate with underlying neurobiology. Nat Commun 15:7987.

39. Lotter LD, Dukart J (2024) NiSpace: Neuroimaging Spatial Colocalization Environment. Available at: https://zenodo.org/records/12514623 [Accessed July 15, 2026].

40. Markello RD, Hansen JY, Liu Z-Q, Bazinet V, Shafiei G, Suárez LE, Blostein N, Seidlitz J, Baillet S, Satterthwaite TD, Chakravarty MM, Raznahan A, Misic B (2022) neuromaps: structural and functional interpretation of brain maps. Nat Methods 19:1472–1479.

41. Markello RD, Spreng RN, Luh W-M, Anderson AK, De Rosa E (2018) Segregation of the human basal forebrain using resting state functional MRI. NeuroImage 173:287–297.

42. Mesulam M-M, Geula C (1988) Nucleus basalis (Ch4) and cortical cholinergic innervation in the human brain: Observations based on the distribution of acetylcholinesterase and choline acetyltransferase. J Comp Neurol 275:216–240.

43. Mesulam M-M, Geula C, Bothwell MA, Hersh LB (1989) Human reticular formation: Cholinergic neurons of the pedunculopontine and laterodorsal tegmental nuclei and some cytochemical comparisons to forebrain cholinergic neurons. J Comp Neurol 283:611–633.

44. Mesulam M-M, Mufson EJ, Levey AI, Wainer BH (1983a) Cholinergic innervation of cortex by the basal forebrain: Cytochemistry and cortical connections of the septal area, diagonal band nuclei, nucleus basalis (Substantia innominata), and hypothalamus in the rhesus monkey. J Comp Neurol 214:170–197.

45. Mesulam M-M, Mufson EJ, Wainer BH, Levey AI (1983b) Central cholinergic pathways in the rat: An overview based on an alternative nomenclature (Ch1–Ch6). Neuroscience 10:1185–1201.

46. Mieling M, Meier H, Bunzeck N (2023) Structural degeneration of the nucleus basalis of Meynert in mild cognitive impairment and Alzheimer’s disease – Evidence from an MRI-based meta-analysis. Neurosci Biobehav Rev 154:105393.

47. Mogg AJ, Eessalu T, Johnson M, Wright R, Sanger HE, Xiao H, Crabtree MG, Smith A, Colvin EM, Schober D, Gehlert D, Jesudason C, Goldsmith PJ, Johnson MP, Felder CC, Barth VN, Broad LM (2018) In Vitro Pharmacological Characterization and In Vivo Validation of LSN3172176 a Novel M1 Selective Muscarinic Receptor Agonist Tracer Molecule for Positron Emission Tomography. J Pharmacol Exp Ther 365:602–613.

48. Mufson EJ, Ginsberg SD, Ikonomovic MD, DeKosky ST (2003) Human cholinergic basal forebrain: chemoanatomy and neurologic dysfunction. J Chem Neuroanat 26:233–242.

49. Muth K, Schönmeyer R, Matura S, Haenschel C, Schröder J, Pantel J (2010) Mild Cognitive Impairment in the Elderly is Associated with Volume Loss of the Cholinergic Basal Forebrain Region. Biol Psychiatry 67:588–591.

50. Naganawa M, Nabulsi N, Henry S, Matuskey D, Lin S-F, Slieker L, Schwarz AJ, Kant N, Jesudason C, Ruley K, Navarro A, Gao H, Ropchan J, Labaree D, Carson RE, Huang Y (2021) First-in-Human Assessment of 11C-LSN3172176, an M1 Muscarinic Acetylcholine Receptor PET Radiotracer. J Nucl Med 62:553–560.

51. Oda Y, Nakanishi I (2000) The distribution of cholinergic neurons in the human central nervous system. Histol Histopathol 15:825–834.

52. Petrou M, Frey KA, Kilbourn MR, Scott PJH, Raffel DM, Bohnen NI, Müller MLTM, Albin RL, Koeppe RA (2014) In Vivo Imaging of Human Cholinergic Nerve Terminals with (–)-5-18F-Fluoroethoxybenzovesamicol: Biodistribution, Dosimetry, and Tracer Kinetic Analyses. J Nucl Med 55:396–404.

53. Qiu T, Hong H, Zeng Q, Luo X, Wang X, Xu X, Xie F, Li X, Li K, Huang P, Dai S, Zhang M (2023) Degeneration of cholinergic white matter pathways and nucleus basalis of Meynert in individuals with objective subtle cognitive impairment. Neurobiol Aging 132:198–208.

54. Ransmayr G, Cervera P, Hirsch E, Ruberg M, Hersh LB, Duyckaerts C, Hauw J-J, Delumeau C, Agid Y (1989) Choline acetyltransferase-like immunoreactivity in the hippocampal formation of control subjects and patients with Alzheimer’s disease. Neuroscience 32:701–714.

55. Rau A, Philipsen L, Frings L, Müller-Glaw P, Reisert M, Mast H, Sajonz BEA, Jost WH, Urbach H, Weiller C, Hosp JA, Bormann T, Rijntjes M, Hellwig S, Schröter N (2025) Hippocampus and basal forebrain degeneration differentially impact cognition in Lewy body spectrum disorders. Brain 148:2772–2784.

56. Robertson RT, Gallardo KA, Claytor KJ, Ha DH, Ku KH, Yu BP, Lauterborn JC, Wiley RG, Yu J, Gall CM, Leslie FM (1998) Neonatal treatment with 192 IgG-saporin produces long-term forebrain cholinergic deficits and reduces dendritic branching and spine density of neocortical pyramidal neurons. Cereb Cortex 8:142–155.

57. Roqué PJ, Barria A, Zhang X, Hashimoto JG, Costa LG, Guizzetti M (2023) Synaptogenesis by Cholinergic Stimulation of Astrocytes. Neurochem Res 48:3212–3227.

58. Sabri O et al. (2015) First-in-human PET quantification study of cerebral α4β2* nicotinic acetylcholine receptors using the novel specific radioligand (−)-[18F]Flubatine. NeuroImage 118:199–208.

59. Salas R, Sturm R, Boulter J, Biasi MD (2009) Nicotinic Receptors in the Habenulo-Interpeduncular System Are Necessary for Nicotine Withdrawal in Mice. J Neurosci 29:3014–3018.

60. Schaefer A, Kong R, Gordon EM, Laumann TO, Zuo X-N, Holmes AJ, Eickhoff SB, Yeo BTT (2018) Local-Global Parcellation of the Human Cerebral Cortex from Intrinsic Functional Connectivity MRI. Cereb Cortex 28:3095–3114.

61. Scheef L, Grothe MJ, Koppara A, Daamen M, Boecker H, Biersack H, Schild HH, Wagner M, Teipel S, Jessen F (2019) Subregional volume reduction of the cholinergic forebrain in subjective cognitive decline (SCD). NeuroImage Clin 21:101612.

62. Schmitz TW, Mur M, Aghourian M, Bedard M-A, Spreng RN (2018) Longitudinal Alzheimer’s Degeneration Reflects the Spatial Topography of Cholinergic Basal Forebrain Projections. Cell Rep 24:38–46.

63. Schmitz TW, Spreng RN (2016) Basal forebrain degeneration precedes and predicts the cortical spread of Alzheimer’s pathology. Nat Commun 7:13249.

64. Scholtens LH, de Reus MA, de Lange SC, Schmidt R, van den Heuvel MP (2018) An MRI Von Economo – Koskinas atlas. NeuroImage 170:249–256.

65. Schulz J, Brandl F, Grothe MJ, Kirschner M, Kaiser S, Schmidt A, Borgwardt S, Priller J, Sorg C, Avram M (2023) Basal-Forebrain Cholinergic Nuclei Alterations are Associated With Medication and Cognitive Deficits Across the Schizophrenia Spectrum. Schizophr Bull 49:1530–1541.

66. Schumacher J, Taylor J-P, Hamilton CA, Firbank M, Cromarty RA, Donaghy PC, Roberts G, Allan L, Lloyd J, Durcan R, Barnett N, O’Brien JT, Thomas AJ (2021) In vivo nucleus basalis of Meynert degeneration in mild cognitive impairment with Lewy bodies. NeuroImage Clin 30:102604.

67. Selden NR, Gitelman DR, Salamon-Murayama N, Parrish TB, Mesulam MM (1998) Trajectories of cholinergic pathways within the cerebral hemispheres of the human brain. Brain 121:2249–2257.

68. Shafiee N, Fonov V, Dadar M, Spreng RN, Collins DL (2024) Degeneration in Nucleus basalis of Meynert signals earliest stage of Alzheimer’s disease progression. Neurobiol Aging 139:54–63.

69. Sharma PK, Gentleman S, Dexter DT, Pienaar IS (2025) Stereological analysis of cholinergic neurons within bilateral pedunculopontine nuclei in health and when affected by Parkinson’s disease. Brain Pathol 35:e70011.

70. Sĭmić G, Mrzljak L, Fučić A, Winblad B, Lovrić H, Kostović I (1999) Nucleus subputaminalis (ayala): the still disregarded magnocellular component of the basal forebrain may be human specific and connected with the cortical speech area. Neuroscience 89:73–89.

71. Simpraga S, Alvarez-Jimenez R, Mansvelder HD, van Gerven JMA, Groeneveld GJ, Poil S-S, Linkenkaer-Hansen K (2017) EEG machine learning for accurate detection of cholinergic intervention and Alzheimer’s disease. Sci Rep 7:5775.

72. Simpraga S, Mansvelder HD, Groeneveld GJ, Prins S, Hart EP, Poil S-S, Linkenkaer-Hansen K (2018) An EEG nicotinic acetylcholine index to assess the efficacy of pro-cognitive compounds. Clin Neurophysiol 129:2325–2332.

73. Slater NM, Melzer TR, Myall DJ, Anderson TJ, Dalrymple-Alford JC (2024) Cholinergic Basal Forebrain Integrity and Cognition in Parkinson’s Disease: A Reappraisal of Magnetic Resonance Imaging Evidence. Mov Disord 39:2155–2172.

74. Teipel SJ et al. (2018) Basal Forebrain Volume, but Not Hippocampal Volume, Is a Predictor of Global Cognitive Decline in Patients With Alzheimer’s Disease Treated With Cholinesterase Inhibitors. Front Neurol 9:642.

75. Teipel SJ, Flatz W, Ackl N, Grothe M, Kilimann I, Bokde ALW, Grinberg L, Amaro E, Kljajevic V, Alho E, Knels C, Ebert A, Heinsen H, Danek A (2014) Brain atrophy in primary progressive aphasia involves the cholinergic basal forebrain and Ayala’s nucleus. Psychiatry Res Neuroimaging 221:187–194.

76. Teipel SJ, Meindl T, Grinberg L, Grothe M, Cantero JL, Reiser MF, Möller H-J, Heinsen H, Hampel H (2011) The cholinergic system in mild cognitive impairment and Alzheimer’s disease: An in vivo MRI and DTI study. Hum Brain Mapp 32:1349–1362.

77. Tian Y, Margulies DS, Breakspear M, Zalesky A (2020) Topographic organization of the human subcortex unveiled with functional connectivity gradients. Nat Neurosci 23:1421–1432.

78. Van Essen DC et al. (2012) The Human Connectome Project: A data acquisition perspective. NeuroImage 62:2222–2231.

79. Voegtle A, Mohrbutter C, Hils J, Schulz S, Weuthen A, Brämer U, Ullsperger M, Sweeney-Reed CM (2024) Cholinergic modulation of motor sequence learning. Eur J Neurosci 60:3706–3718.

80. Wang Y, Zhan M, Roebroeck A, De Weerd P, Kashyap S, Roberts MJ (2022) Inconsistencies in atlas-based volumetric measures of the human nucleus basalis of Meynert: A need for high-resolution alternatives. NeuroImage 259:119421.

81. Wearn A, Tremblay SA, Tardif CL, Leppert IR, Gauthier CJ, Baracchini G, Hughes C, Hewan P, Tremblay-Mercier J, Rosa-Neto P, Poirier J, Villeneuve S, Schmitz TW, Turner GR, Spreng RN (2024) Neuromodulatory subcortical nucleus integrity is associated with white matter microstructure, tauopathy and APOE status. Nat Commun 15:4706.

82. Weuthen A, Kirschner H, Ullsperger M (2025) Error-driven upregulation of memory representations. Commun Psychol 3:17.

83. Whitehouse P, Price D, Struble R, Clark A, Coyle J, DeLong MR (1982) Alzheimer’s Disease and Senile Dementia: Loss of Neurons in the Basal Forebrain. Science 215:1237–1239.

84. Zaborszky L, Hoemke L, Mohlberg H, Schleicher A, Amunts K, Zilles K (2008) Stereotaxic probabilistic maps of the magnocellular cell groups in human basal forebrain. NeuroImage 42:1127–1141.

